# Biophysical validation of explainable AI for functional brain imaging: bridging cellular mechanisms and network dynamics

**DOI:** 10.1101/2025.02.24.639808

**Authors:** Anthony Strock, Trang-Anh E. Nghiem, Alex De Lecea, Atif Hassan, Srikanth Ryali, Vinod Menon

## Abstract

Deep neural networks have revolutionized functional neuroimaging analysis but remain “black boxes,” concealing which brain mechanisms and regions drive their predictions—a critical limitation for clinical neuroscience. Here we develop and validate an explainable AI (xAI) framework to test whether feature attribution techniques can reliably recover brain regions affected by excitation/inhibition (E/I) imbalance—a fundamental dysregulation implicated in autism, schizophrenia, and other neuropsychiatric disorders. We employed complementary simulation approaches: recurrent neural networks for controlled parameter exploration, and The Virtual Brain simulator incorporating empirically-derived human and mouse connectomes to model E/I balance alterations with unprecedented biological realism. Through systematic validation, we demonstrate that Integrated Gradients and DeepLIFT methods reliably identify brain regions affected by E/I imbalance across challenging conditions, including high noise, low prevalence, and subtle neurophysiological alterations. This performance remains robust across species and anatomical scales, from 68-region human to 426-region mouse connectomes. Application to the ABIDE autism dataset (N=834) reveal convergence between our biophysically grounded simulations and empirical findings, providing computational support for E/I imbalance mechanisms in autism. This work establishes essential tools and data for interpretation of deep learning models in functional neuroimaging, and enables hypothesis-driven analysis of cellular mechanisms across neuropsychiatric disorders.

## Introduction

Deep neural networks (DNNs) are increasingly central to human brain research, offering powerful tools for scientific discovery and clinical applications (1, 2, 3, 4, 5). These models have demonstrated remarkable success in identifying patterns and making predictions from complex neuroimaging data (3, 4, 5). However, a fundamental challenge persists: DNN models typically function as “black boxes” offering no insight into the neurobiological and cellular mechanisms underlying their performance. This lack of interpretability poses a significant obstacle in neuroscience, where identifying meaningful biomarkers is essential for advancing our understanding of brain function and dysfunction. Here we develop and validate an explainable AI (xAI) framework for interpreting deep neural networks in functional neuroimaging, enabling hypothesis-driven testing of specific cellular mechanisms.

Excitation/inhibition (E/I) imbalance has emerged as a fundamental framework for understanding various neurological and psychiatric disorders, including autism and schizophrenia (6, 7, 8, 9). This dysregulation manifests through altered glutaminergic and GABAergic signalling and regional changes in neurotransmitter concentrations, as demonstrated by converging evidence from cellular studies and magnetic resonance spectroscopy (9, 10, 11). In autism specifically, disrupted E/I balance appears to be a key mechanism underlying its core symptoms of restricted social interests and circumscribed behaviors (12, 13), with evidence spanning multiple scales of investigation from cellular studies showing altered GABAergic signalling in animal models, to human neuroimaging revealing regional GABA concentration changes (14, 15). These E/I alterations are hypothesized to specifically affect nodes of the default mode network (16), which plays a central role in self-referential processing, social cognition, and autobiographical memory retrieval - cognitive functions notably impaired in autism, schizophrenia and many other psychiatric and neurological disorders (16, 17, 18).

However, identifying the specific brain regions or networks where E/I imbalance occurs remains a critical challenge due to methodological limitations. Current methods to measure imbalance, such as magnetic resonance spectroscopy, require targeting specific hypothesized areas a priori. This limitation has crucial consequences in clinical settings, where brain regional targets must be identified for interventions altering E/I imbalance through localized brain stimulation. Therefore, providing data-driven, testable hypotheses for E/I imbalance localization could play a central role in uncovering targets for experimental and therapeutic protocols.

Functional brain imaging offers a promising approach to probe circuit function and dysfunction associated with E/I imbalance. Conventional functional MRI analyses rely heavily on pre-engineered features, focusing primarily on static inter-regional connectivity measures or predefined activation patterns (19, 20, 21). Recent attempts to incorporate neural networks, including graph-based approaches (22, 23) and surface-based convolutional networks (20), still largely reduce temporal dynamics to static representations. However, E/I imbalance mechanisms manifest through dynamic alterations in neural signalling that may only be detectable when preserving the full spatiotemporal complexity of brain activity patterns (24, 25, 26).

Deep neural networks that analyze dynamic fMRI time series data without feature engineering offer a promising solution to this challenge. Spatiotemporal DNNs (stDNNs), in particular, address limitations of traditional approaches by learning discriminative features directly from raw temporal signals, modelling a hierarchy of temporal scales preserving the full complexity of brain activity patterns (3, 4, 27). These models have shown promise in detecting aberrant circuit dynamics in psychiatric disorders where E/I imbalance is prominent, including autism and psychosis (3, 4). Previous applications have demonstrated that dynamic circuit alterations detected by stDNNs can predict symptom-defined subtypes in autism (4) and genetically-defined subtypes in psychosis (3), with findings showing robustness across multisite datasets (3, 4). However, the critical question remains: can we validate that these models are indeed able to detect E/I imbalance mechanisms?

This validation challenge is particularly acute because xAI methods themselves must be rigorously tested with known ground truth before we can confidently use them to guide mechanistic understanding or treatment strategies. Without such validation, it would not be possible to distinguish between genuine biological insights and algorithmic artifacts. This interpretability gap is particularly problematic for clinical applications, where understanding the biological basis of model decisions is essential for developing targeted interventions (25, 28). This reflects a fundamental tension in clinical machine learning between achieving high predictive accuracy and gaining mechanistic insights that can inform therapeutic development (28, 29).

To address these challenges, we developed and validated an xAI framework using two complementary mechanistic simulation approaches to test whether attribution methods can reliably recover brain regions affected by E/I imbalance. A key goal was to develop recurrent neural network (RNN) and The Virtual Brain (TVB) models that capture essential brain dynamics and E/I balance alterations, enabling systematic testing of whether stDNN-based xAI methods can accurately identify affected regions under controlled conditions with known ground truth – something impossible with real neuroimaging data alone. First, we employ RNNs to generate synthetic fMRI data with precisely controlled parameters. Second, we utilized TVB simulator (30) to create biophysically informed simulations incorporating realistic brain connectivity from both human (31) and mouse connectomes (32). This dual approach enables validation across different scales and species while maintaining known ground truth.

In both approaches, we systematically investigated how various neurophysiological factors – including noise levels, prevalence of brain regions affected by E/I imbalance, and severity of E/I alterations – impact feature attribution in neuroimaging analyses. By manipulating these parameters, we could simulate different scenarios of E/I imbalance, which is crucial for understanding neurological disorders such as autism (13) and schizophrenia (33).

Our study addresses four key research questions that establish validation protocols for neuroimaging xAI. First, can feature attribution methods reliably localize regions with altered E/I balance when clinical differences arise from these cellular-level mechanisms? We test this using carefully constructed RNN simulations with known ground truth, systematically varying the prevalence of affected regions (1 to 10% of nodes), severity of E/I alterations

(δ = 0.1-0.5), and signal-to-noise ratios (−10dB to 10dB). Second, how robust is this localization across different biological scales and anatomical complexity at the whole-brain level? To address this, we extend validation using biophysically realistic TVB simulations incorporating empirically-derived connectomes from both human (68 regions) and mouse (426 regions) brains. Third, how does attribution accuracy depend on method choice and parameter selection? We systematically compare multiple xAI approaches to gradient-based feature attribution— Integrated Gradients (34), DeepLIFT (35), Input X Gradient (36),

Guided Backprop (37)— that are computationally tractable and inexpensive but whose relative performance remains actively contested even in foundational domains like visual image categorization (36, 38). We also examine how baseline choice affects attribution in dynamic brain imaging, a crucial methodological question unique to time series data. Fourth, what is the interpretability of attribution maps when applied to real clinical data? We apply our validated methods to the multisite ABIDE dataset (N = 834)(4), testing whether regions identified by xAI methods align with known neurobiological theories of E/I imbalance in autism, enabling us to assess whether our biophysically grounded validation framework provides sufficient confidence for interpreting deep learning models in clinical neuroimaging.

Our comprehensive framework provides methodological guidelines for xAI applications in clinical neuroimaging and generates testable hypotheses about target sites and mechanisms to guide future experimental and therapeutic efforts.

## Results

### RNN simulations and validation

We first evaluate xAI methods using recurrent neural network (RNN) simulations, which allowed systematic control over key parameters while maintaining temporal dynamics characteristic of fMRI data. These simulations generated synthetic fMRI time series from two groups: one with typical network dynamics and another with hyperexcitable nodes, mimicking localized E/I imbalance.

#### Model design and parameter space

The RNN model generated time series data for 100 regions, with adjustable parameters controlling: signal-to-noise ratio (SNR: −10dB to 10dB), prevalence of affected regions (1 to 10% of nodes), and severity of hyperexcitability (δ: 0.1 to 0.5). Network connectivity was defined using small-world topology to reflect brain-like architecture, while regional dynamics were modelled using nonlinear activation functions convolved with a canonical hemodynamic response (see Methods).

#### Classifier performance

Our stDNN classifier demonstrated robust performance across simulation conditions (**Fig S1**). Even at high noise levels, with 5% prevalence, and small hyperexcitability (δ = 0.1), the classifier achieved perfect discrimination (accuracy = 100%) (−10dB SNR) (**Fig S1A**). Performance remained robust down to 1% prevalence, though convergence required more training epochs (**Fig S1B**). The classifier successfully detected group differences across hyperexcitability levels above δ = 0.1, with faster convergence at higher severities (**Fig S1C**).

#### xAI analysis

IG successfully identified hyperexcited nodes across all simulation conditions (**Fig 2**).To establish a consistent threshold across xAI methods and simulation parameters, we normalized all attributions by dividing it by its maximum value across ROIs. With a normalized attribution threshold of 0.5, IG achieved perfect detection (F1 score = 100%) of affected network nodes even under challenging conditions: high noise (−10dB SNR, **Fig 2A-C**), low prevalence (1%, **Fig 2D-F**), and subtle hyperexcitability (δ = 0.1, **Fig 2G-I**).

**Figure 1.**
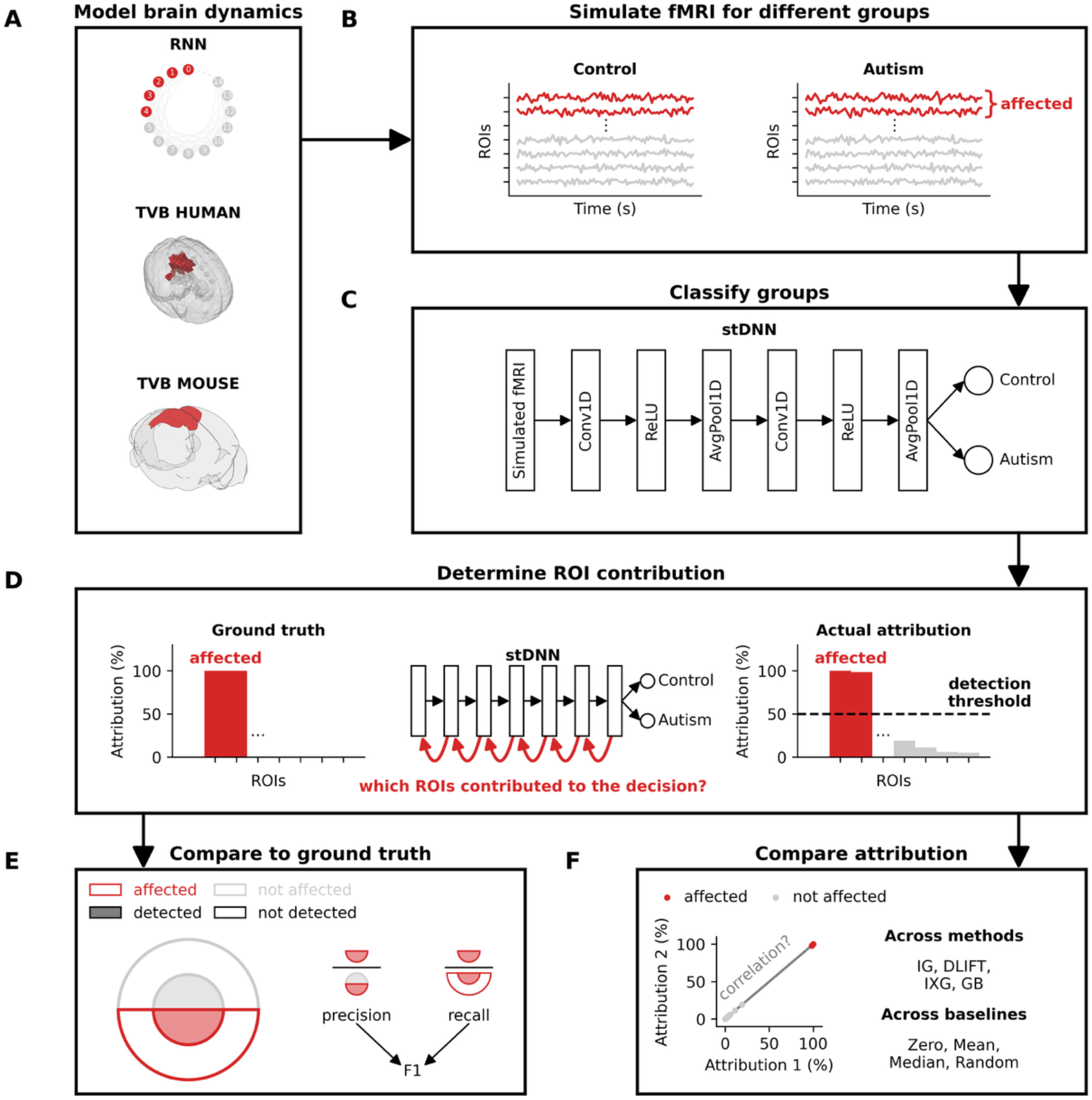
Overview of explainable AI framework for E/I imbalance detection. Simulated multivariate fMRI datasets consisting of two classes are generated using two complementary simulation approaches: (1) recurrent neural networks (RNNs) for controlled parameter exploration, and (2) biophysically realistic whole-brain simulations using The Virtual Brain (TVB) simulator, which incorporates empirically-derived structural connectomes and models neural dynamics through coupled excitatory and inhibitory populations in each brain region. Both simulation approaches generate two classes that share identical structural connectivity (adjacency matrix), but differ in excitation/inhibition balance—with some nodes (highlighted in red) exhibiting altered E/I balance in one population through modified inhibitory conductance parameters. A spatiotemporal deep neural network (stDNN) consisting of 1D convolutional layers is trained to classify the two populations based on their dynamic fMRI signals. Feature attribution methods are then applied to the trained model to identify which brain regions drive classification decisions, testing the hypothesis that these methods can reliably recover the known locations of E/I alterations.

**Figure 2.**
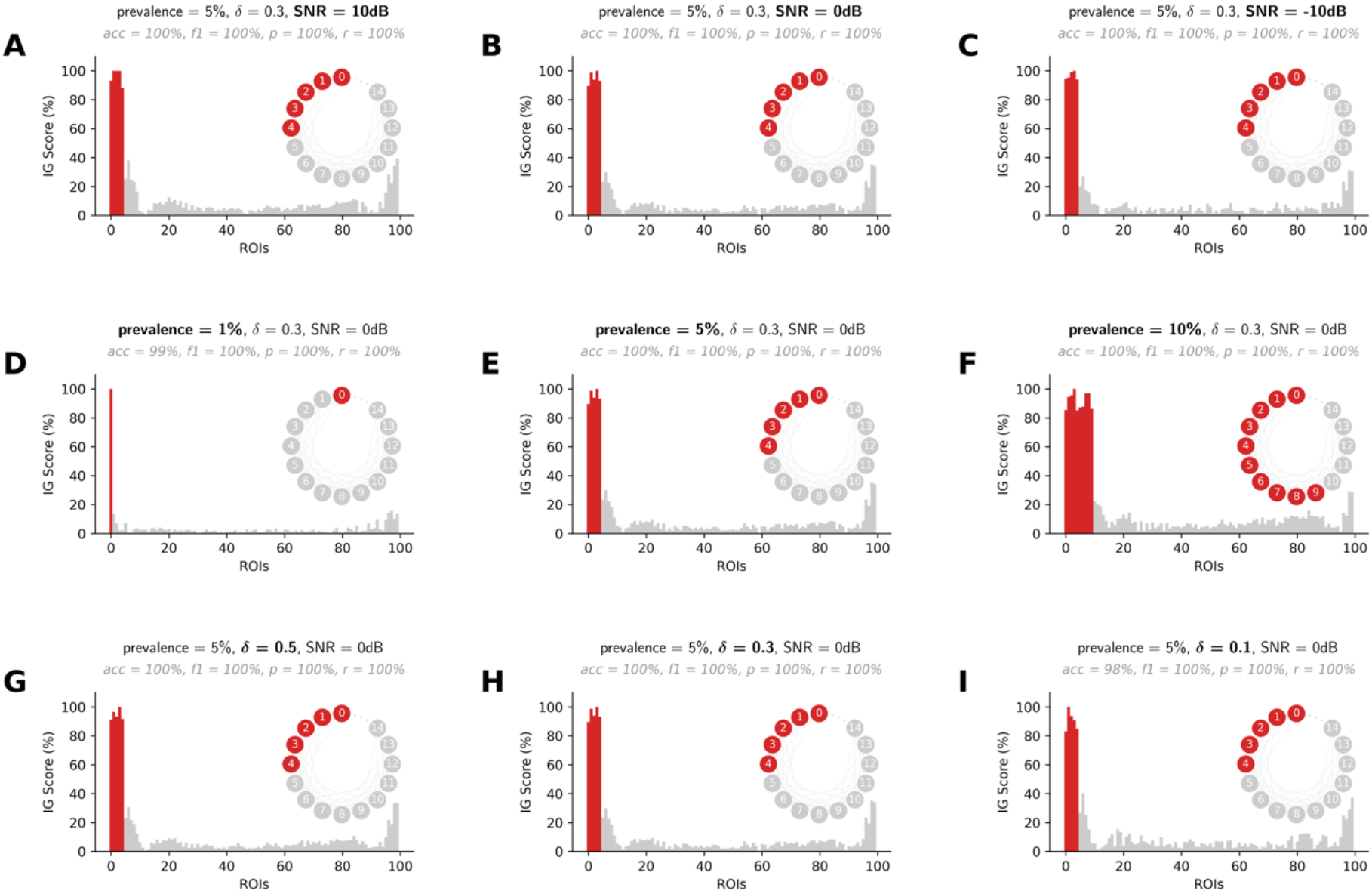
Validation of Integrated Gradients for detecting E/I imbalance in RNN simulations. Performance of Integrated Gradients (IG) in identifying regions with altered excitation/inhibition balance in small-world RNN simulations across varying signal-to-noise ratios (SNR), prevalence rates, and E/I imbalance severity. Regions affected by E/I alterations (highlighted in red) were reliably identified with perfect accuracy (F1-score = 1.0) across all tested conditions. (**A-C**) IG attribution scores for SNRs of 10, 0 and −10 dB with 5% prevalence and severity δ = 0.3. (**D-F**) IG scores for prevalence rates of 1%, 5% and 10% with SNR of 0 dB and severity δ = 0.3. (**G-I**) IG scores for E/I imbalance severity δ = 0.5, δ = 0.3, and δ = 0.2 with SNR of 0 dB and 5% prevalence.

Remarkably, IG’s performance remained stable across these conditions, suggesting robust feature detection even with minimal signal.

These RNN simulation results established that IG can reliably identify affected brain regions across a wide range of simulation conditions.

### Human TVB Simulations

We next evaluated xAI methods using biophysically informed simulations with The Virtual Brain (TVB), incorporating realistic human brain connectivity and dynamics. These simulations provided a more stringent test of our approach while maintaining known ground truth alterations in specific brain regions.

#### Biophysical model implementation

Using structural connectivity from human diffusion imaging (68-region Desikan-Killiany atlas(39)), we simulated whole-brain dynamics with explicit modelling of excitatory and inhibitory populations in each region. We created two groups by introducing E/I imbalance in one group through altered inhibitory synaptic conductance (ΔQi = 0.1-0.5nS): first targeting only bilateral posterior cingulate cortex (PCC), then progressively including precuneus (PCun) and angular gyrus (AnG) to test effects of anatomical extent on classification and IG performance.

#### Classifier performance

The stDNN classifier achieved robust discrimination between groups across simulation conditions (**Fig S2**). For large imbalance in PCC-only, classification accuracy remained above 90% for SNR levels down to −10dB (**Fig S2A**). Convergence time decreased with increasing anatomical extent of E/I imbalance, with fastest classification when alterations included PCC, PCun, and AnG (**Fig S2B**). The classifier showed sensitivity to E/I imbalance severity, with reliable discrimination (accuracy > 90%) for ΔQi ≥ 0.3nS (**Fig S2C**).

#### xAI results

IG successfully identified regions with altered E/I balance across conditions (**Fig 3**). With PCC-only alterations, IG showed high attributions specifically localized in the bilateral PCC across SNR levels (**Fig 3A-C**). When additional regions were affected, IG accurately detected the expanded anatomical extent of alterations (**Fig 3D-F**), maintaining precise localization even as the number of affected regions increased. Attribution patterns remained stable across E/I imbalance severities (**Fig 3G-I**), though stronger alterations (ΔQi = 0.5nS) produced more pronounced attribution scores. Importantly, regions without simulated E/I alterations consistently showed attribution scores below our normalized attribution threshold of 0.5, demonstrating high specificity in feature detection.

**Figure 3.**
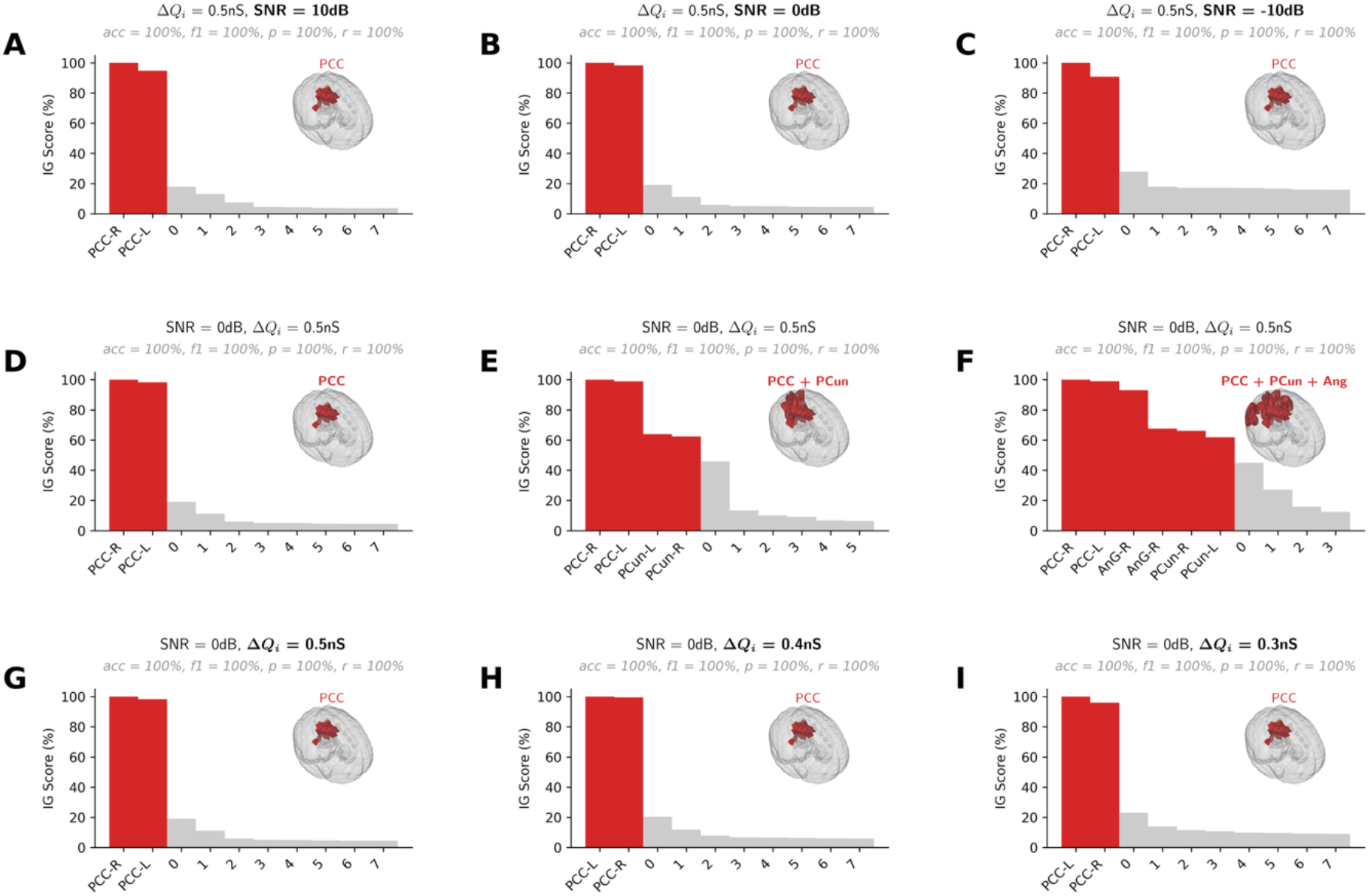
Cross-species validation using human connectome TVB simulations. Performance of Integrated Gradients (IG) in detecting E/I imbalance in biophysically realistic whole-brain simulations using human structural connectivity (68-region Desikan-Killiany atlas). E/I imbalance was modeled through altered inhibitory conductance (ΔQi) in default mode network regions. Affected regions (highlighted in red) were consistently identified across all conditions. (**A-C**) IG scores for SNRs of 10, 0 and −10 dB with E/I imbalance isolated to bilateral posterior cingulate cortex (PCC), ΔQi = 0.5nS. (**D-F**) IG scores for progressive anatomical extent: PCC only, PCC+precuneus (PCun), and PCC+PCun+angular gyrus (AnG) with ΔQi = 0.5nS and SNR of 0 dB. (**G-I**) IG scores for varying E/I imbalance severity in PCC: ΔQi = 0.5nS, 0.4nS, and 0.3nS with SNR of 0 dB.

These TVB simulation results demonstrate that xAI methods can reliably identify biologically realistic E/I alterations in the context of whole-brain dynamics and authentic human brain connectivity. The successful detection of affected regions across varying anatomical distributions and E/I imbalance severities supports the method’s potential utility for identifying circuit alterations in clinical populations.

### Mouse TVB simulations

To test the generalizability of our approach across species and scales, we conducted parallel analyses using the mouse connectome. Unlike human diffusion imaging-based connectivity, the Allen Brain Atlas provides invasively validated, directional connectivity data across 426 regions, offering higher spatial resolution and more direct measurement of anatomical connections.

#### Model implementation

We adapted our TVB simulations to the mouse brain, maintaining the same biophysical parameters for excitatory and inhibitory populations while incorporating mouse-specific structural connectivity. E/I imbalance was introduced in the retrosplenial cortex (RSC), the key DMN hub in rodents analogous to the human PCC. We systematically varied inhibitory conductance (ΔQi: 0.1 to 0.5nS) to parallel our human simulations.

#### Classifier performance

The stDNN classifier showed robust but distinct performance patterns with mouse data (**Fig S3**). Classification accuracy remained above 90% for moderate noise levels (0dB SNR) with RSC-only alterations (**Fig S3A**). Performance improved when E/I imbalance extended to additional regions (RSC+Cg, RSC+Cg+PrL; **Fig S3B**), though requiring more training epochs than for human simulations. The classifier maintained sensitivity to E/I imbalance severity, achieving reliable discrimination for ΔQi ≥ 0.3nS (**Fig S3C**).

#### xAI results

IG successfully identified affected regions of the mouse brain, though with some species-specific patterns (**Fig 4**). With RSC-only alterations, IG showed enhanced attributions precisely localized in all RSC subregions only for SNR levels down to 0dB (**Fig 4A-C**). When E/I imbalance extended to additional regions, IG accurately detected the broader anatomical distribution (**Fig 4D-F**), maintaining precision and recall above 80%. Hierarchical attribution strengths, varying across subregions, were observed here due to more structurally distinct regions affected here than in human TVB simulations. This pattern held across E/I imbalance severities (**Fig 4G-I**), suggesting robust detection of regional hierarchies in the mouse brain.

**Figure 4.**
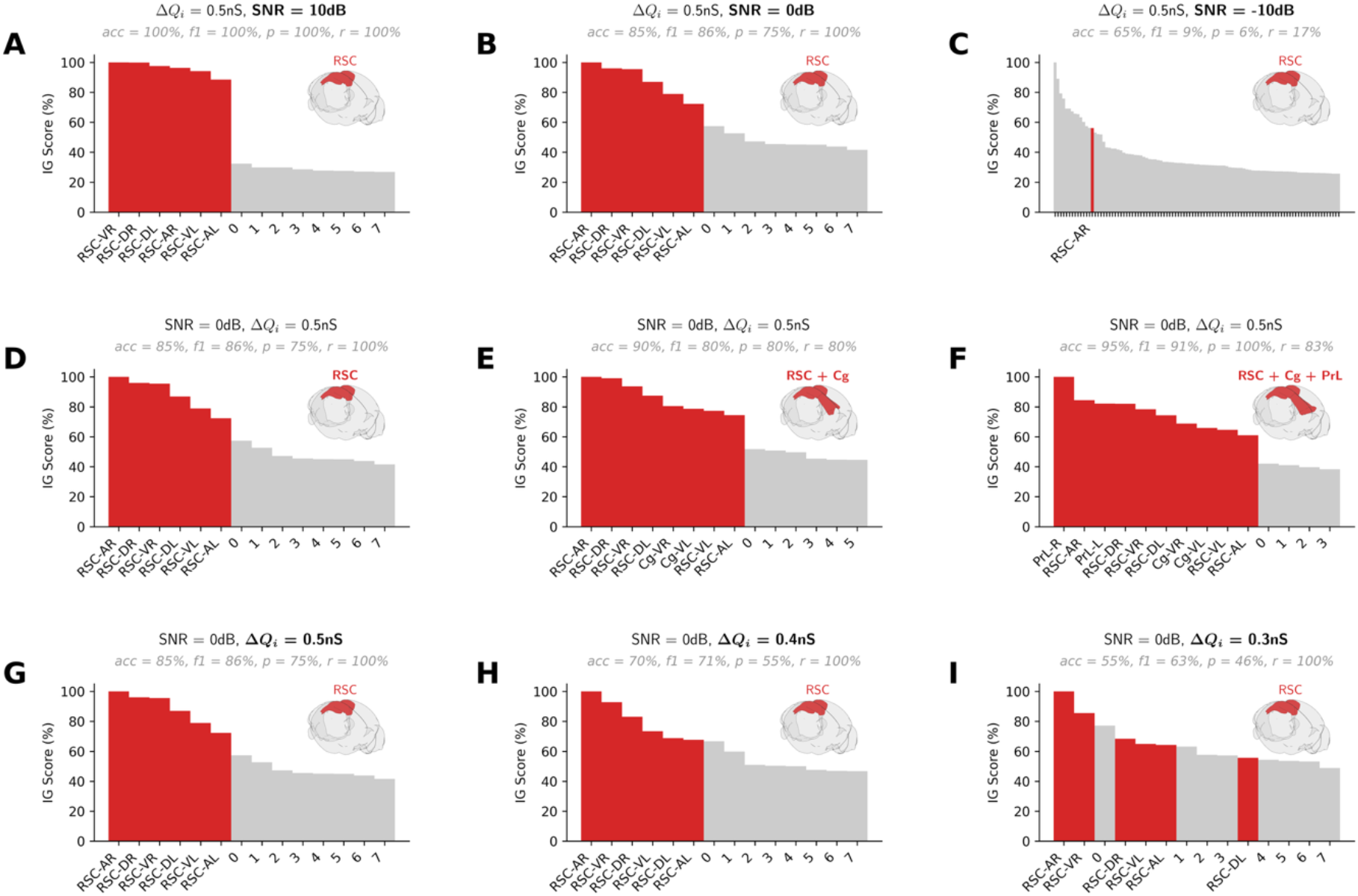
Cross-species validation using mouse connectome TVB simulations. Performance of Integrated Gradients (IG) in detecting E/I imbalance in high-resolution mouse brain simulations using Allen Brain Atlas connectivity (426 regions). E/I imbalance was introduced in retrosplenial cortex (RSC), the rodent analog of human posterior cingulate cortex. (**A-C**) IG scores for SNRs of 10, 0 and −10 dB with E/I imbalance in RSC, ΔQi = 0.5nS. (**D-F**) IG scores for progressive anatomical extent: RSC only, RSC+cingulate cortex (Cg), and RSC+Cg+prelimbic cortex (PrL) with ΔQi = 0.5nS and SNR of 0 dB. (**G-I**) IG scores for varying E/I imbalance severity in RSC: ΔQi = 0.5nS, 0.4nS, and 0.3nS with SNR of 0 dB.

The successful cross-species validation, particularly the detection of analogous regions (RSC/PCC) despite different scales and connectivity patterns, demonstrates the robustness of our xAI approach. The higher spatial resolution of the mouse connectome suggests the potential relevance of the stDNN and IG approach to high-dimensional data and to reveal additional insights about subregional specificity while maintaining reliable detection of E/I alterations.

### ABIDE fMRI dataset analysis

After validating our approach using simulated data, we applied these methods to the Autism Brain Imaging Data Exchange (ABIDE) dataset to identify brain regions that reliably distinguish individuals with autism from neurotypical controls. This multi-site dataset (N=834; 419 autism, 415 controls) provided a robust test of our methods in real-world clinical data.

#### Classifier performance

As reported previously (4), our stDNN achieved significant classification accuracy across the ABIDE dataset (78.2 ± 2.84%, using 5-fold cross validation), reliably distinguishing autism from control participants. Performance remained stable across different site splits and cross-validation folds, demonstrating robustness to data collection variability. Notably, classification accuracy was maintained even with highly heterogeneous symptomatology across autistic individuals and multi-site imaging data (4).

#### xAI analysis

xAI analysis revealed a consistent set of discriminative brain regions (**Fig 5**). IG identified the bilateral posterior cingulate cortex (PCC) and right precuneus as the regions most strongly contributing to classification, with attribution scores above our normalized attribution threshold of 0.5. This finding was remarkably stable across multiple analyses as described below.

**Figure 5.**
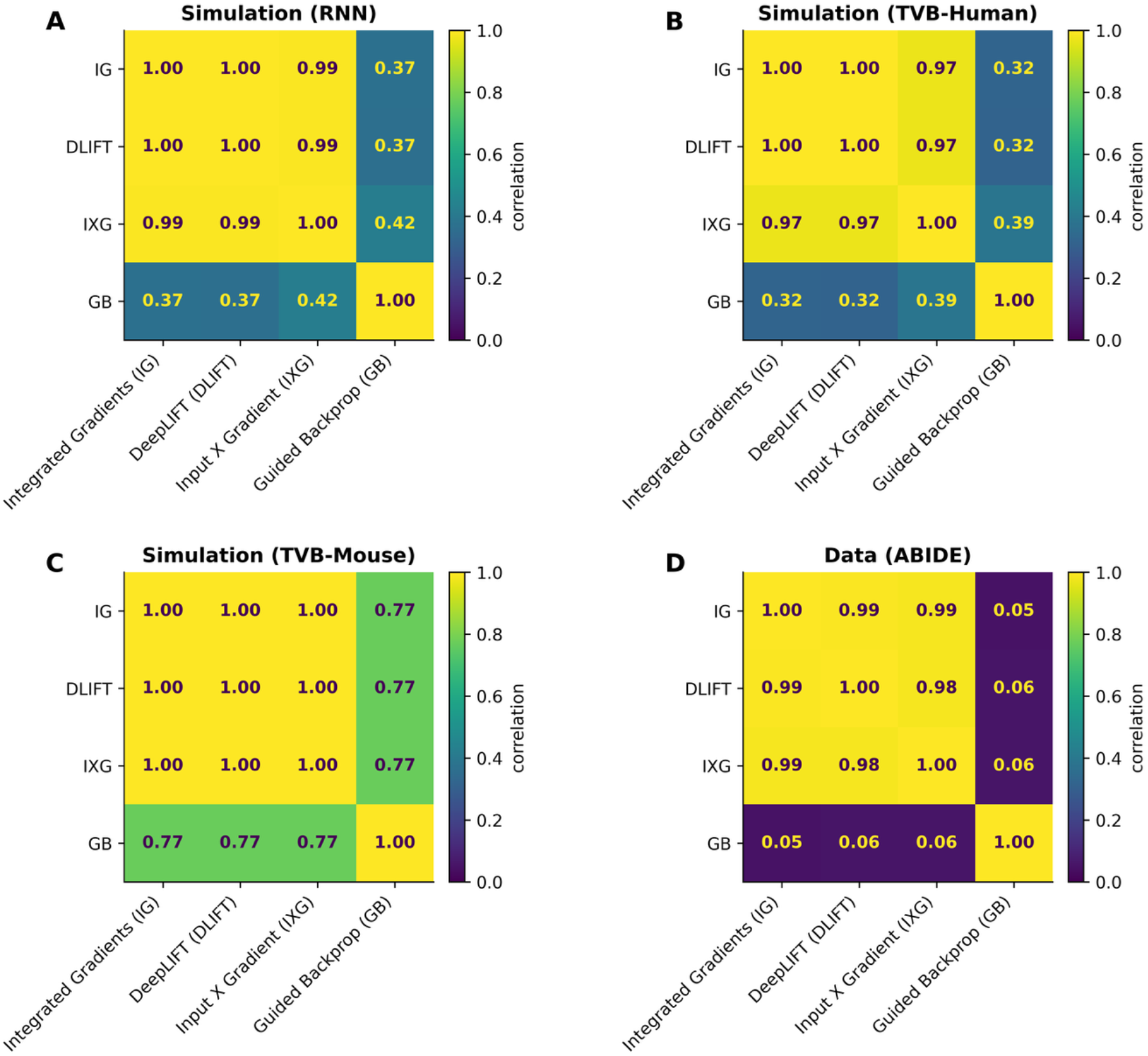
Application to autism neuroimaging data reveals default mode network involvement. Feature attribution scores from the ABIDE autism dataset (N = 834) using validated xAI methods. (**A**) Integrated Gradients (IG), (**B**) DeepLIFT, and (**C**) Input X Gradient (IXG) show highly consistent attribution patterns, with bilateral posterior cingulate cortex (PCC) and right precuneus (PCun) exceeding the validated threshold of 0.5. (**D**) Guided Backpropagation (GB) shows diffuse, less discriminative attributions across many regions, with lower scores for PCC and precuneus.

The convergence between our empirical findings and simulation results provides computational support for the role of E/I imbalance in autism, particularly within the DMN. The identification of these regions is consistent with previous studies showing altered DMN function in autism and adds mechanistic insight by suggesting these alterations may reflect underlying E/I imbalance.

### Comparison of xAI methods

To establish the reliability of our approach, we systematically compared multiple xAI methods across both simulated and empirical data. We evaluated four methods: Integrated Gradients (IG), DeepLIFT, Input X Gradient (IXG), and Guided Backpropagation (GB), each representing different theoretical approaches to explaining model decisions.

#### RNN simulations

In our controlled RNN simulations, IG, DeepLIFT, and IXG showed remarkably consistent performance (**Fig 6A**). These methods demonstrated high correlations in their attribution patterns (r > 0.99) and achieved similar accuracy as measured by F1score in identifying hyperexcitable nodes (**Fig S4A**). In contrast, GB showed weaker correlations with other methods (r ≤ 0.42) and impaired F1 scores (**Fig S4A**) due to reduced precision (**Fig S4B**) but not recall (**Fig S4C**) in detecting affected regions.

**Figure 6.**
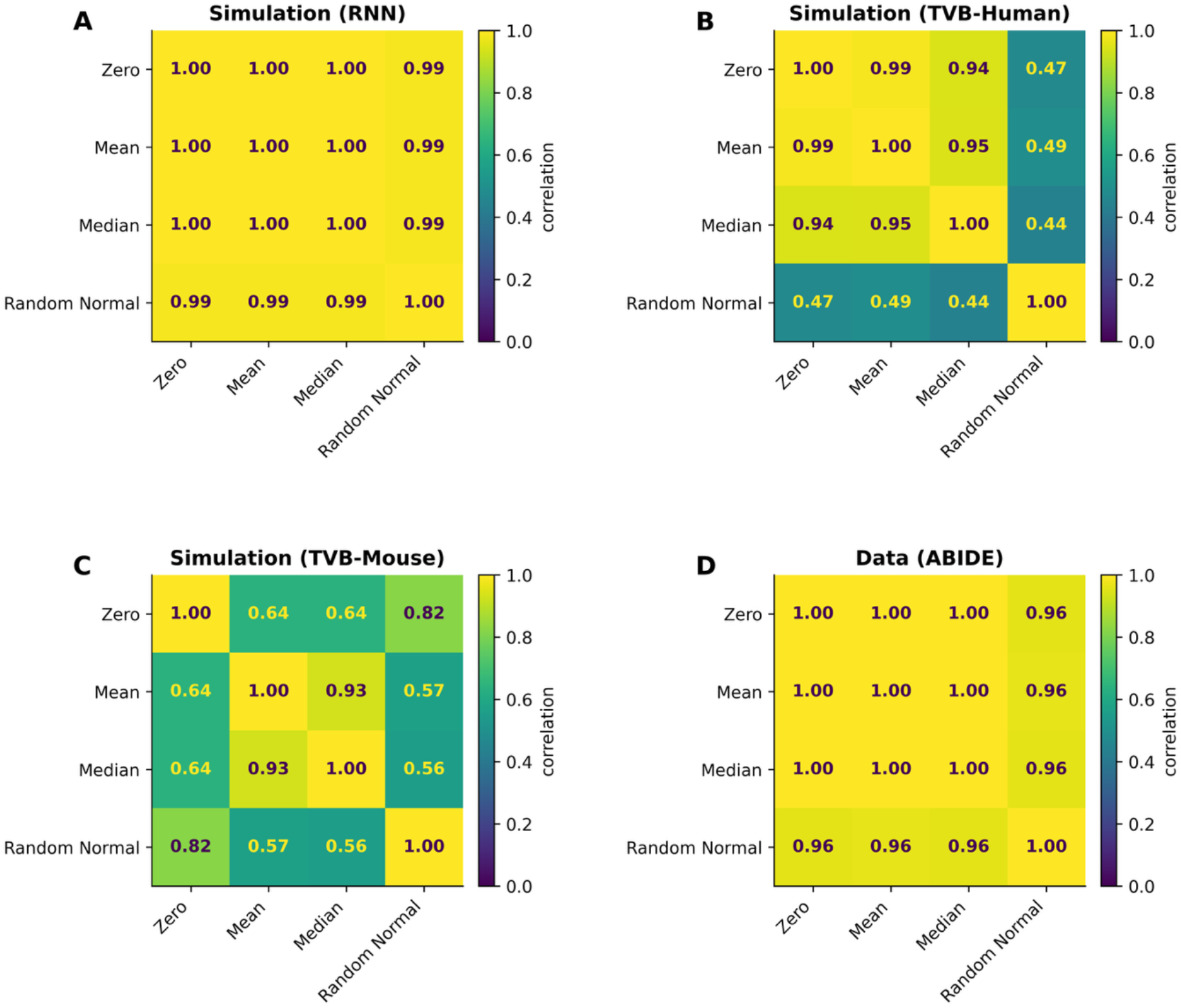
Systematic comparison of feature attribution methods across validation conditions. Correlation matrices showing agreement between different attribution methods across simulation and empirical data. (**A-D**) Pairwise correlations between attribution patterns from Integrated Gradients (IG), DeepLIFT (DL), Input X Gradient (IXG), and Guided Backpropagation (GB) in: (**A**) RNN simulations, (**B**) human TVB simulations, (**C**) mouse TVB simulations, and (**D**) ABIDE autism data. High correlations (r > 0.95) between IG, DeepLIFT, and IXG demonstrate method consistency, while Guided Backpropagation shows divergent patterns across all conditions.

#### Human TVB simulations

In human TVB data, IG, DeepLIFT, and IXG maintained strong correlations (r > 0.97, **Fig 6B**) in identified regions with affected E/I balance. Weaker correlations were seen between attribution profiles of GB and other methods (r ≤ 0.39). Poorer F1 scores (**Fig S4D**) at identifying affected nodes due to lower precision (**Fig S4E**) but not recall (**Fig S4F**) was also observed for GB compared to other methods.

#### Mouse TVB simulations

Similar patterns emerged in mouse TVB simulations, where these methods showed high agreement (r > 0.99, **Fig 6C**) despite the increased anatomical complexity of the mouse connectome. In contrast, GB showed weaker correlations with other methods (r = 0.77) and reduced F1 score due to low precision in detecting affected regions (**Fig S4G-I**).

#### ABIDE fMRI data

In the ABIDE dataset, IG, DeepLIFT, and IXG identified highly consistent sets of discriminative regions (r > 0.98, **Fig 6D**). All three methods highlighted the bilateral PCC and right precuneus as key regions distinguishing autism from controls. GB showed markedly different attribution patterns (r < 0.06), suggesting potential limitations in its clinical applications.

These results demonstrate that IG, DeepLIFT, and IXG provide consistent and reliable feature attribution across different experimental conditions and scales of brain organization, while GB shows divergent patterns and reduced accuracy, suggesting it may be less suitable for neuroimaging applications.

### Impact of baseline choice in Integrated Gradients

The choice of baseline represents a critical methodological consideration when applying Integrated Gradients to neuroimaging data. Unlike static image analysis where a black (zero) image serves as a natural baseline, the optimal reference point for dynamic brain data is not immediately obvious. We therefore systematically evaluated multiple baseline choices including the zero, mean, median across the ROIs of the input timeseries, and random across our simulation frameworks and empirical data.

#### RNN simulations

In RNN simulations, zero, mean, median, and random baselines showed remarkably high correlations in their attribution patterns (r ≥ 0.99, **Fig 7A**) and achieved equivalently good performance in identifying affected regions (**Fig S5A-C**).

**Figure 7.**
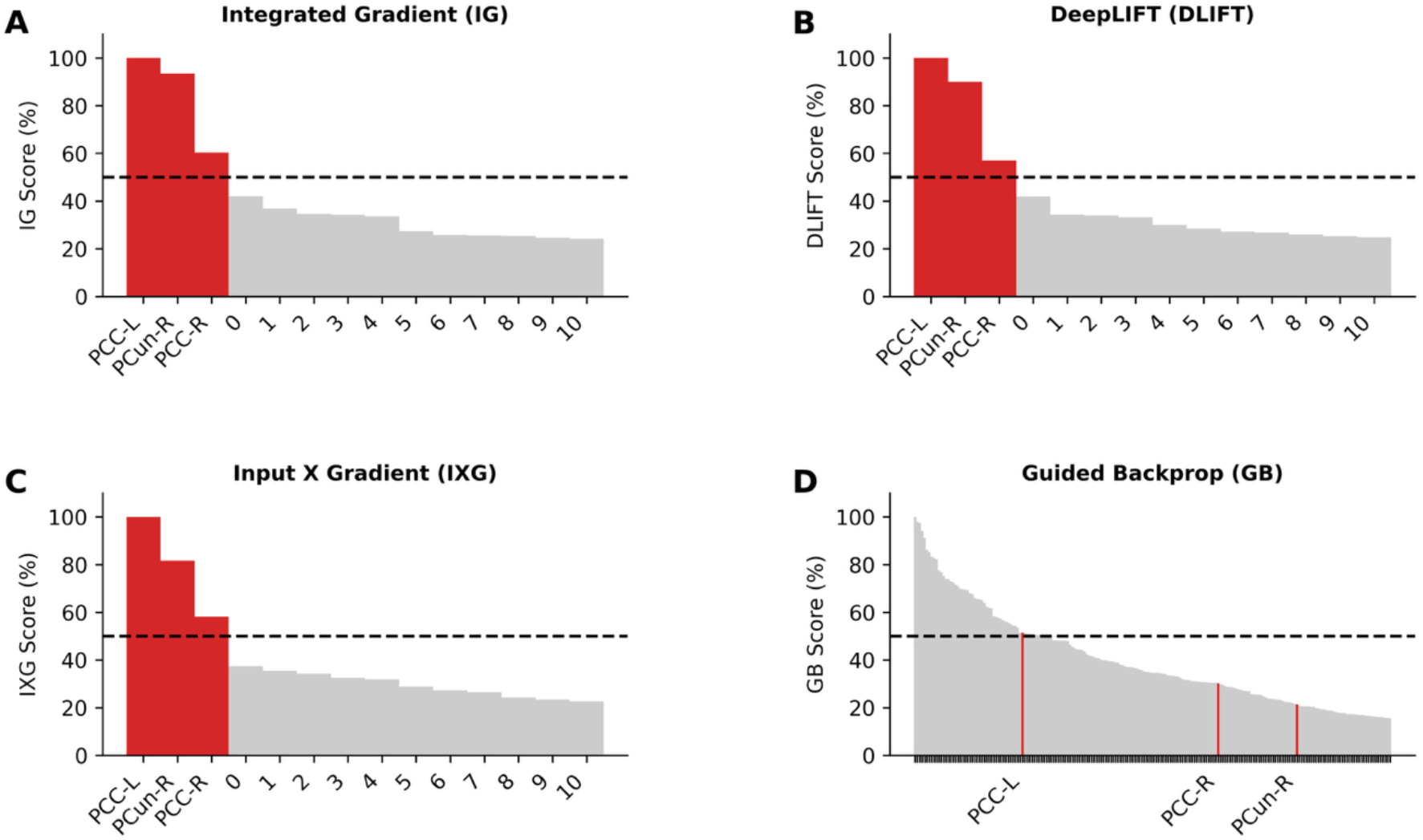
Robustness of Integrated Gradients to baseline selection in dynamic brain imaging. Correlation matrices evaluating the impact of baseline choice on IG attribution patterns. (**A-D**) Pairwise correlations between attribution maps generated using median, mean, zero, and random noise baselines in: (**A**) RNN simulations, (**B**) human TVB simulations, (**C**) mouse TVB simulations, and (**D**) ABIDE data. High correlations between median, mean, and zero baselines (r > 0.94) demonstrate robustness, while random baselines show lower consistency, indicating they should be avoided in neuroimaging applications.

#### Human TVB simulations

This consistency extended to the more complex TVB simulations, where these three baselines maintained strong correlations despite added biological complexity (r ≥ 0.94 for human, **Fig 7B**). The preservation of attribution patterns across baseline choices suggests robust feature detection rather than methodological artifacts. In contrast, random baselines showed weaker correlations with other methods (r < 0.49) and reduced F1 scores in detecting affected regions (**Fig S5D**) due to both impaired precision (**Fig S5E**) and recall (**Fig S5F**) compared to other baseline choices.

#### Mouse TVB simulations

On higher-dimensional mouse TVB simulations yielded high correlations between mean and median baselines (r = 0.93) as well as between zero and random normal baselines (r = 0.82). Weaker correlations (r ≤ 0.64) were found between other baseline pairs (**Fig 7C**). As with RNN simulations, no significant differences across baseline choices were found as to their accuracy scores at identifying affected regions (**Fig S5G-I**).

#### ABIDE fMRI data

Baseline robustness was further confirmed in empirical data, with zero, mean, and median baselines showing highly correlated attribution patterns in the ABIDE dataset (r > 0.99, **Fig 7D**). Importantly, random baselines consistently showed poorer performance across all conditions, with significantly lower correlations with other baselines (typically r ≤ 0.96).

These results demonstrate that zero, mean, and median baselines all provide stable and consistent feature attribution in neuroimaging applications, but random baselines should be avoided due to their poor reproducibility.

## Discussion

DNN models have emerged as powerful tools for analyzing functional MRI data, offering unprecedented accuracy in predicting cognitive states and clinical outcomes (3, 4). However, their inherent complexity and “black box” nature have limited our ability to determine which neuroanatomical features and neurophysiological mechanisms drive their predictions. This opacity poses a significant challenge in neuroscience, where interpretability is crucial for advancing our understanding of brain function and dysfunction and for developing targeted therapeutic interventions. The core challenge is that without ground-truth validation, we cannot assess whether xAI attribution methods identify genuine neurobiological mechanisms or algorithmic artifacts. This validation gap has severely limited our understanding of xAI attributions, particularly for clinical applications where identifying the biological basis of model decisions is essential for guiding mechanistic hypotheses and treatment development.

Here we addressed this fundamental challenge by developing and validating an xAI framework to test whether attribution methods can reliably recover brain regions affected by excitation/inhibition (E/I) imbalance, a well-established cellular mechanism implicated across multiple neuropsychiatric disorders (6, 7, 8, 9). A key innovation was creating biophysically realistic simulations using RNN and TVB models that capture essential brain dynamics and E/I balance alterations with unprecedented biological realism, enabling systematic hypothesis testing under controlled conditions with known ground truth – something impossible with real neuroimaging data alone. Our findings demonstrate that xAI methods can reliably identify brain regions affected by E/I imbalance across a wide range of conditions establishing the first validated framework for biological interpretation of deep learning models in functional neuroimaging.

### Feature attribution methods reliably localize regions with altered E/I balance

RNN simulations provided the first systematic answer to whether feature attribution methods can reliably localize regions with pathophysiology by generating synthetic fMRI data with precisely controlled E/I alterations. We demonstrated that Integrated Gradients can reliably identify affected brain regions across challenging conditions that mirror real-world neuroimaging scenarios (**Fig 2**). The robustness across extreme conditions is particularly important for clinical applications, where signal quality varies significantly between individuals and disease-related alterations may be subtle or spatially sparse. Our systematic validation across prevalence rates (1 to 10% of nodes), severity levels (δ: 0.1 to 0.5), and noise conditions (SNR: −10dB to 10dB) provides performance benchmarks that enable researchers to assess the reliability of xAI results in their own datasets. The robustness across extreme conditions is particularly important for clinical applications, where signal quality varies significantly between individuals and disease-related alterations may be subtle or spatially sparse.

Our results demonstrate that xAI methods can reliably bridge the gap between cellular-level mechanisms and network-level patterns, making xAI findings more interpretable from a cellular mechanism standpoint in that xAI can localize E/I imbalance.

### Robustness of E/I imbalance localization across different biological scales and anatomical complexity

A critical question for any neuroimaging method is whether findings generalize across different brain organizations, spatial scales, and species. Validation must account for substantial differences in connectivity patterns, anatomical resolution, and measurement approaches between human and animal models. Our TVB simulations incorporating empirically-derived human and mouse connectomes demonstrated remarkable robustness across biological scales (**Figs 3 and 4**). Despite substantial differences in brain parcellation, connectivity patterns, and measurement approaches (DTI-based vs viral tracing), xAI methods successfully identified altered E/I balance in analogous brain regions—the posterior cingulate cortex in humans and retrosplenial cortex in mice.

The human TVB simulations, incorporating structural connectivity from the Human Connectome Project (31), allowed us to model E/I imbalance within realistic brain network architectures. IG successfully identified regions with altered inhibitory conductance across different anatomical extents and severity levels, maintaining precise localization even as the number of affected regions increased. The mouse simulations provided validation at higher spatial resolution using the Allen Brain Atlas’s experimentally validated connectivity data (32), demonstrating that our approach scales effectively to finer anatomical detail.

This cross-species validation represents a significant advance by providing computational support for translational research using mouse models to study E/I imbalance mechanisms relevant to human disorders. The consistency between species despite different scales and connectivity patterns demonstrates the robustness of our xAI approach and suggests broad applicability across neuroimaging applications. This finding is particularly important as it establishes that xAI methods detect biological mechanisms rather than artifacts of specific connectome properties, enabling translation between preclinical models and human studies.

### Robustness of xAI methods choice and parameter selection on attribution accuracy

A critical unresolved question in explainable AI has been whether different attribution methods yield consistent and biologically meaningful results. Even in foundational domains like visual image categorization, the relative performance of these methods remains actively contested despite their widespread use (36, 38). For neuroimaging, this uncertainty is particularly problematic because inconsistent results across methods would undermine confidence in any biological interpretation. Additionally, methodological parameters like baseline selection pose unique challenges for dynamic functional brain imaging that have not been systematically addressed.

Our systematic comparison of four prominent xAI methods – Integrated Gradients (34), DeepLIFT (DL) (35), Input X Gradient (IXG) (36), Guided Backprop (GB) (37) – revealed striking patterns of consistency and divergence (**Fig 5**). IG, DeepLIFT, and IXG showed remarkably high correlations (r > 0.95) across all simulation conditions, from simple RNN models to complex TVB simulations with both human and mouse connectomes. This consistency suggests these methods capture fundamental features of network dynamics rather than methodological artifacts. In contrast, Guided Backpropagation showed notably weaker correlations with other methods (r < 0.4 in human data, r < 0.77 in mouse data) and reduced accuracy in identifying ground truth affected regions.

Regarding parameter selection, we addressed the critical question of baseline choice for dynamic brain imaging (**Fig 6**). Unlike static image analysis where zero baselines are natural, temporally complex fMRI signals require careful consideration of what constitutes “absence of signal.” Our systematic evaluation revealed that zero, mean, and median baselines yield highly correlated attribution patterns (r > 0.94) across simulation conditions and real data, providing methodological flexibility while maintaining interpretive validity. However, random baselines showed poor reproducibility and should be avoided.

These results provide the first systematic guidelines for method and parameter selection in neuroimaging xAI. The high consistency between IG, DeepLIFT, and IXG establishes that reliable biological interpretation is achievable when appropriate methods are selected, providing methodological clarity and validated protocols for future studies seeking to uncover mechanistic alterations in autism and other neuropsychiatric disorders.

### Clinical implications and future applications

Our validated xAI framework enables several important clinical applications and testable hypotheses. Most directly, our findings E/I imbalance disruptions can be manifest as perturbations in xAI attribution maps. This interpretation can be empirically tested by targeting magnetic resonance spectroscopy measurements to quantify GABA concentrations in regions identified by feature attribution in individual patients’ neuroimaging data.

The demonstrated ability of xAI methods to reliably localize E/I imbalance across different severity levels and anatomical distributions opens new possibilities for clinical stratification. Our framework could potentially identify subgroups of individuals with autism based on the regional patterns and severity of E/I disruptions revealed through attribution maps. This approach is particularly relevant given the substantial heterogeneity in autism symptom profiles and severity, which may require different treatment strategies depending on underlying neurobiological subtypes.

However, several important limitations must be acknowledged before clinical implementation. Our validation was performed on a single disorder and dataset, requiring replication across independent cohorts and multiple neuropsychiatric conditions. Additionally, the relationship between xAI attributions and actual GABA concentrations measured through spectroscopy remains to be empirically established. Future studies should directly correlate attribution scores with biochemical measurements and clinical outcomes to validate the biological relevance of these computational findings.

Beyond autism, our validation framework can be extended to other disorders where E/I imbalance is hypothesized, such as schizophrenia and epilepsy. The biophysical simulation approach can also be adapted to test other cellular mechanisms including neuromodulation and synaptic plasticity, enabling systematic validation of xAI methods across different neurobiological hypotheses. This represents a general framework for hypothesis-driven interpretation of deep learning models in clinical neuroscience.

## Conclusion

We developed and validated a biophysically realistic simulation framework for testing whether explainable AI methods can reliably identify cellular-level mechanisms underlying neuropsychiatric disorders. Our systematic validation demonstrates that Integrated Gradients and DeepLIFT methods can accurately localize regions affected by excitation/inhibition imbalance, maintain robust performance across species and anatomical scales, and translate meaningfully to clinical populations.The scientific significance of this work extends beyond methodological validation. By establishing that explainable AI methods can reliably recover known E/I alterations in controlled simulations, we provide the neuroimaging community with essential tools for confident biological interpretation of deep learning models. This work represents a foundation for interpretable and biologically informed applications of artificial intelligence across neuroimaging research.

## Materials and Methods

We evaluate the accuracy of the relevant inputs retrieved by xAI methods using two different simulation models for synthesizing fMRI timeseries. The first simulation model is a simple recurrent neural network model where the differences between two classes is simulated using different functional connectivity between the brain regions as well as the local excitatory/inhibitory (E/I) imbalances in some subset of brain regions. The primary purpose of this simple model is to investigate the performance of the xAI methods as a function of prevalence rates and SNRs and to find the optimal thresholds on the xAI scores to separate the most relevant brain features from the irrelevant ones. The second model is a more realistic biophysical model for autism which is implemented using The Virtual Brain stimulator. The differences between the two classes is created by introducing differences in the local E/I imbalances in some set of brain regions. These regions are chosen based on our recent findings (4, 40) in the autism population. We simulate timeseries of nodes (or ROIs) consisting of two classes which differ either with respect to the connectivity of ROIs in the two classes or differ with respect to the local excitability of some chosen nodes. We then train a classifier to predict the class label given the timeseries of the nodes. Finally, we examine how each xAI method discovers important nodes or features in each class. We hypothesize that these xAI methods would correctly identify the nodes that differ in functional connectivity or and their local excitability between the classes.

### Simulated fMRI data with a recurrent neural network model

We simulated the whole-brain fMRI timeseries by the time series of 100 nodes or regions of interest (ROIs). To generate simulated fMRI data of two classes, we assumed that the difference between the two classes lies in their local excitability on some chosen nodes. We assumed that the network consists of 100 nodes defined using an adjacency matrices *A* of a randomly sampled connected Watts–Strogatz small-world graphs (41), each of the 100 nodes joined with its *k* = 10 nearest neighbors in a ring topology with a probability *p* = 0.1 of rewiring. The local level of excitability for the two classes, *W*_0_ and *W*_1_, are defined as *W*_*k*_ = ***Λ***_*k*_*A*^*T*^, where 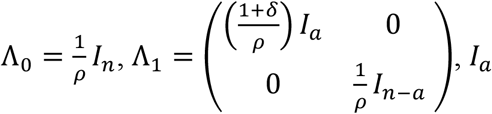 and *I*_*n*–*a*_ are respectively the *a* × *a* and the (*n* − *a*) × (*n* − *a*) the identity matrices, *n* = 100 is the total number of nodes, *a* is the number of nodes affected by the difference in local excitability, *ρ* is the spectral radius of the matrix *A*, and *δ* > 0 measures the amplitude of the difference between the two classes.

We simulate several fMRI datasets with varying degrees of differences in their local excitability. We quantify these differences using (1) a prevalence rate that describes the ratio of the number of nodes that differ in local excitability to the overall number of nodes (i.e. the ratio 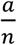), and (2) the severity of the hyper-excitability defined as the ratio of local difference in excitability in affected nodes to the level of excitability (i.e. the ratio 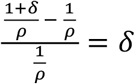). We hypothesize that the difficulty of finding the discriminating nodes increases when the prevalence rate or the hyper-excitability level drops.

We simulated fMRI data using a Recurrent Neural Network (RNN) convolved with a canonical hemodynamic response function (HRF), which can be described with the following dynamical system:

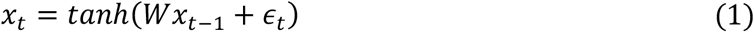

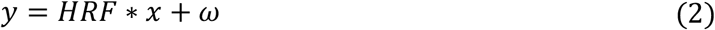

Where *W* ∈ {*W*_0_, *W*_1_} is the recurrent weight matrix, *y*_*t*_ represents the BOLD fMRI signal of multiple regions of interests (ROIs) at time *t, x*_*t*_ represents hidden neural variables giving rise to the BOLD signal at time *t, ϵ*_*t*_ = *N*(0, *σ*_*ϵ*_) represents the intrinsic neural noise, *ω*_*t*_ = *N*(0, *σ*_*ω*_) represents the measurement noise which is independent across regions, *HRF* is a canonical hemodynamic response function with a sampling period of 1s (42).

We chose *σ*_*ω*_ to achieve some fixed level of Signal to Noise Ratio (*SNR*) as follows:

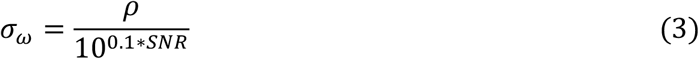

where *σ*_*ω*_ is the variance of the noise and *ρ* is the maximal variance of the signal across ROIs that we empirically computed after simulating *x*_*t*_ dynamics. An example of this simulated data for each of the two classes over time can be seen in **Fig 1**.

We investigated the performance of various xAI methods with the prevalence rates ranging from 1 to 10% in steps of 1%, the hyper-excitability amplitude ranging from 0.0 to 0.5 in steps of 0.1, and the *SNR* ranging from −10dB to 10dB in steps of 10dB. For each set of parameters, 400 random samples were generated, with 80% used for training and 20% held out for testing.

### Simulated fMRI data with The Virtual Brain Simulator

We used a more realistic biophysical model to generate synthetic fMRI datasets for two classes representing the neurotypical and autism populations to investigate the performance of the xAI methods. Specifically, we simulated fMRI data using a model of whole-brain dynamics where biophysical parameters can be varied at the regional level. The model was implemented in The Virtual Brain (TVB) simulator (30, 43) in previous work (44). Briefly, each simulated region comprises two neural populations, one of excitatory (E) and one of inhibitory (I) neurons, whose mean firing rates across E and I populations are described by the following differential equations:

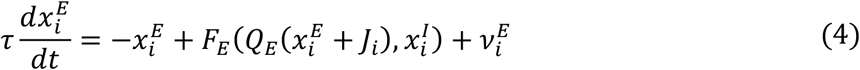

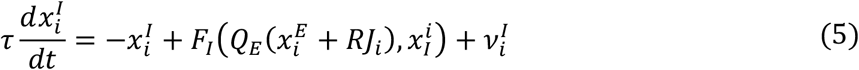

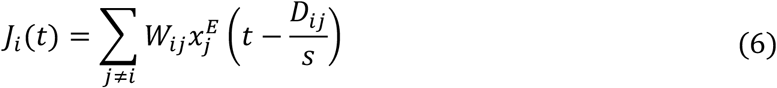

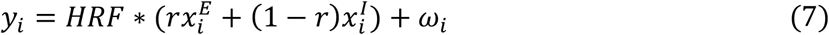

Where 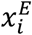 and 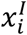 are respectively the firing rates of excitatory and inhibitory populations in region *i, J*_*i*_ represents the input into region *i* of other regions long-range excitatory connections between regions, *R* = 1.4 is the ratio between the conductance of long-range excitatory-to-excitatory and excitatory-to-inhibitory synapses, *W*and *D* characterize respectively the number and the length of white-matter fibres between regions between regions both empirically derived from synaptic tracing in the mouse brain, *s* = 3*m*/*s* represents the speed of transmission between regions, *Q*_*E*_ = 1 *nS* and *Q*_*I*_ represents respectively the synaptic strength of excitation and inhibition, *v*_*E*_ and *v*_*I*_ are standard normal integration noise, the transfer functions *F*_*E*_ and *F*_*I*_ which relates each population’s output rate to the input it receives from itself and other populations were fitted to reproduce the input-output relations of integrate-and-fire neurons in spiking simulations (45), *y* represents the BOLD fMRI signal of multiple regions of interests (ROIs), *ω*_*i*_ = *N*(0, *σ*_*ω*_) represents the measurement noise in region *i, HRF* is a canonical hemodynamic response function. As previously we chose *σ*_*ω*_ to achieve some fixed level of Signal to Noise Ratio (*SNR*) as follows:

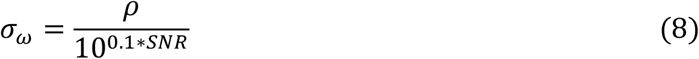

where *σ*_*ω*_ is the variance of the noise and *ρ* is the maximal variance of the signal across ROIs that we empirically compute after simulation.

For the human brain, tract weights and lengths were obtained with diffusion tensor imaging of a healthy adult’s brain (44) (data publicly available at https://zenodo.org/records/4263723). The connectome was partitioned according to the commonly used 68-ROI Desikan-Killiany-Tourville atlas (46). For the mouse brain, the structural connectome was taken from a tracer-derived structural connectome of the mouse brain from the Allen Brain Institute (32) with a 426-ROI parcellation. Inter-regional synaptic delays were modelled based on the matrix of tract lengths from the same source (32). Synaptic weights and delays were taken to be the same from the left and right brain hemispheres.

The datasets for the autism class was generated by introducing E/I imbalance in a subset of ROIs. Specifically, the biophysical parameter *Q*_*I*_ capturing inhibitory synaptic conductance was modified specifically in regions of the default mode network where excitation/inhibition imbalance has been observed in humans and rodent models of autism and hypothesized to underlie symptoms. For the human brain, *Q*_*I*_ was altered bilaterally progressively in the posterior cingulate cortex, in the precuneus (PCun) and in the angular gyrus (AnG). In the mouse brain, *Q*_*I*_ was locally modified progressively in the bilateral agranular, ventral, and dorsal subdivisions of the retrosplenial cortex (RSC), in the cingulate cortex (Cg) and in the prelimbic cortex (PrL).

We simulated time series of 100 s for all ROIs for each connectome. This procedure was repeated with 100 different random seeds that affected initial conditions and random noise realizations, 80 seeds were used for training and 20 held out for testing. To simulate fMRI data, the firing rate time series were then convolved with a canonical hemodynamic response function with TR of 1s. To test the robustness of the xAI methods across different noise levels, in both human and mouse models, measurement noise was added to the ROI timeseries generated using the above simulation models. To quantify the impacts of severity of E/I imbalance, inhibitory conductance in affected regions was varied between 4.5 nS and 4.9 nS to obtain severity varying from ΔQi = 0.1nS to ΔQi = 0.5nS between affected and unaffected regions. To investigate the effects of prevalence, imbalance was implemented in different subsets of DMN regions of varying size – in the human model, we compared imbalance bilaterally in the PCC only, in the PCC and PCun only, and in the PCC, PCun and AnG, while in the mouse model, we compared imbalance bilaterally in the RSC only, the RSC and Cg only, and the RSC, Cg, and PrL.

### ABIDE dataset

We leveraged neuroimaging and phenotypic data from the Autism Brain Imaging Data Exchange (ABIDE; http://fcon_1000.projects.nitrc.org/indi/abide/) (1, 2). The dataset consists 419 subjects with autism and 415 typically developing subjects from 14 sites. We used the same preprocessed resting-state fMRI dataset as in previous work (4), excluding the same subject identified as having missing or poor-quality imaging data. The Brainnetome Atlas was applied, comprising 246 ROIs included in the analysis.

### stDNN classifier model for the simulated fMRI datasets and for the ABIDE dataset

We used a two-layer 1D convolutional neural network model for classifying the two groups both simulated fMRI datasets and in the ABIDE dataset, integrating spatial ROI information at each convolutional layer through summation across channels. For simulated fMRI datasets, as shown in **Fig 1C**, our stDNN processes simulated fMRI time series with a 1D convolution, followed by a ReLU nonlinearity and average pooling. It then transforms the features with a second 1D convolution and ReLU, followed by adaptive average pooling that reduces each feature to a single time step. The output is flattened and passed through a linear layer for classification. For ABIDE dataset, following original paper (4), our stDNN processes fMRI time series with two successive 1D convolution–ReLU–max-pooling blocks, averages features across time, and applies dropout. The resulting representation is concatenated with an auxiliary covariate encoding, then passed through a fully connected layer and sigmoid activation for binary classification. We used 80% of the data to train the model, the remaining 20% for testing the model. The code for the model implemented in PyTorch is available at https://github.com/scsnl/Strock_Nghiem_bioRxiv_2025.

### XAI methods

We obtained the discriminating features from each simulated subject by using a xAI method which determines how much each nodes or ROI contributes to the decision made by our classifier for this subject. The simulated fMRI data being temporal, each of these xAI methods provides us with a value for each ROI and for each time step of simulation. We obtain the influence of an ROI by taking the absolute value of the median of the xAI across time step. To evaluate the performance of the attribution method we normalize the attribution by dividing it by the maximum threshold across ROI, and compare ROI with attribution above 0.5 with ROI affected E/I imbalance using standard recall, precision and F1 metrics.

We first evaluated the performance of Integrated Gradient (IG) (34) on the simulated datasets. We examined how IG performance may depend on the choice of baseline which is required for evaluating the path integral for the feature attribution. It is not clear what an optimal baseline is for the neuroimaging applications. Therefore, we investigated several baselines including zero, random, mean, and median of all fMRI time series across the subjects. We finally compared the performance of other gradient based xAI methods such as DeepLIFT (DL) (35), Input X Gradient (IXG) (36), Guided Backprop (GB) (37) implemented in the Captum python library.

### Integrated Gradients

IG directly uses the gradient of the classifier with respect to the input ROIs to define the xAI. This method can be used for any already trained DNN model with a single backpropagation pass. It integrates this gradient over a path in the fMRI input space going to the subject’s simulated fMRI data from a baseline of the same dimension. Formally, IG attribution method can be written as:

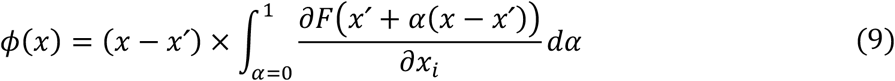

where *x* is the original input, *x*^′^ is the chosen baseline from which we start our integration, *F* represents the function associating fMRI input to its clinical category, and *ϕ* is the function giving the xAI score for a given node x. IG overcomes the problems associated with the simple gradient based approaches such as the noisy gradients and gradient saturation. As IG requires a baseline, we compared the xAI obtained for various baselines: (1) the median of the signal across nodes and samples, (2) the mean of the signal across nodes and samples, (3) the default all-zero baseline, and (4) a random sample drawn from a normal distribution centred about zero with the variance reflecting the variance of the input.

### Deep learning important features (DeepLIFT)

As IG, DeepLIFT is xAI method that can applied to any pretrained DNN model with a single backpropagation pass. This method computes the importance of each input in the prediction of the output. This importance for each input is computed in terms of the difference between the output for a reference or the baseline and the given input. As in the IG, the baseline needs to be chosen based on the domain. It is developed to address the issues with the basic gradient based xAI approaches. As in the case of IG, we evaluate the performance of DeepLIFT for all the baselines as defined above.

### Input X Gradient (IXB)

IXB is also a gradient based attribution method used for obtaining xAIs. This is a very basic method where gradient of the input with respect to the target is computed and then the gradient is multiplied with the input to obtain the xAI scores. If both the gradient of the input and also its magnitude is high then that feature’s attribution is high. Both IG and DeepLIFT methods are refinements over this approach.

### Guided Backpropagation (GB)

This method is developed for explaining the decisions taken by a CNN model by computing the gradient of the output with respect to the input. This method requires a modification of the gradient in the ReLU layers where the gradient to these layers are made zero if the input to the ReLU during the forward pass is negative. This approach is applicable to only the convolutional networks.

### Statistical analysis

Relationships between variables were quantified using Pearson’s correlation coefficient (*r*), and group differences were assessed using two-tailed *t*-tests. All statistical tests were implemented in Python using the SciPy library, with significance evaluated at P < 0.05 (*), P < 0.01 (**), or P < 0.001 (***).

## Data Availability

This study used only existing, publicly available neuroimaging data from the Autism Brain Imaging Data Exchange (ABIDE), hosted on the International Neuroimaging Data-Sharing Initiative (INDI) platform (http://fcon_1000.projects.nitrc.org/indi/abide/). The data were collected and shared by the ABIDE consortium; no new data were collected by the authors. All analyses were purely computational. The code used for the computational model and analyses is available at https://github.com/scsnl/Strock_Nghiem_bioRxiv_2025.

## Supplementary Figures

**Figure S1.**
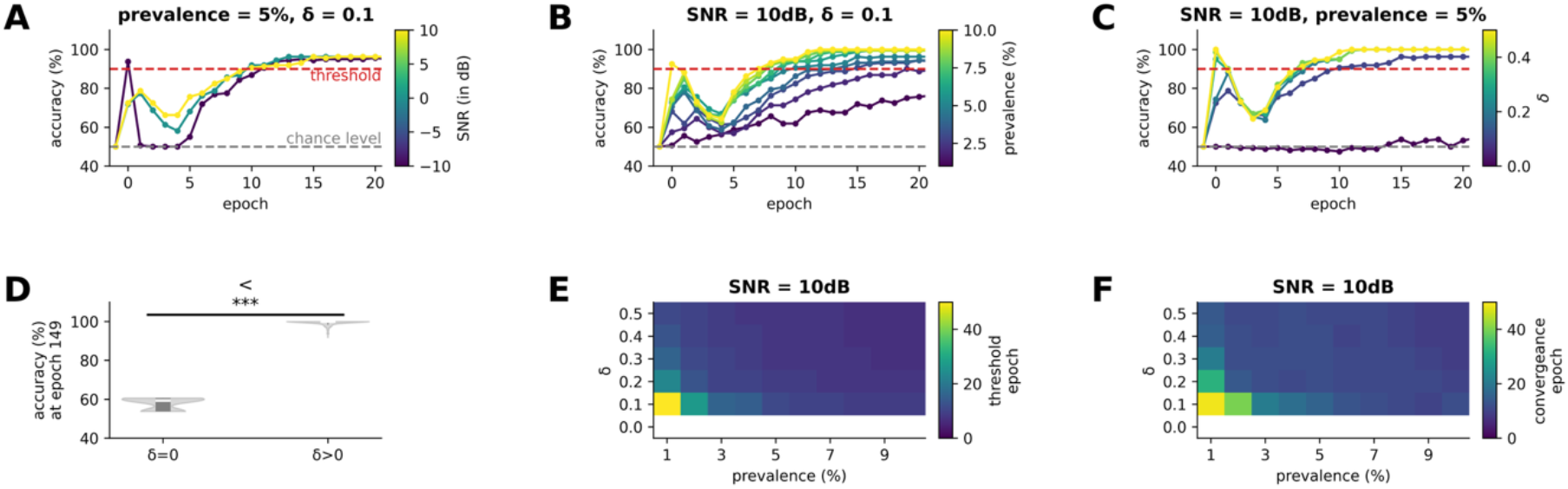
Training and classification performance on simulated data generated using RNNs. (**A-C**) Classification accuracy as a function of training epochs (**A**) for SNRs ranging from −10 dB to 10 dB with a fixed prevalence rate of 5% and hyper-excitability severity of δ = 0.1, (**B**) for prevalence rates ranging from 1 to 10% with fixed SNR of −10 dB hyper-excitability severity of δ=0.1, and (**C**) for hyper-excitability severity varying from δ = 0.0 to δ = 0.5 with fixed SNR of −10 dB and prevalence rate of 5%. (**D**) Classification accuracy at epoch 149 for hyper-excitability severity of δ = 0.0 to δ > 0.0. (**E-F**) Training epoch during which a 90% accuracy threshold is passed (**E**) and convergence epoch (**F**) as a function of prevalence rate and hyper-excitability severity for an SNR of 10 dB. *** P < 0.001.

**Figure S2.**
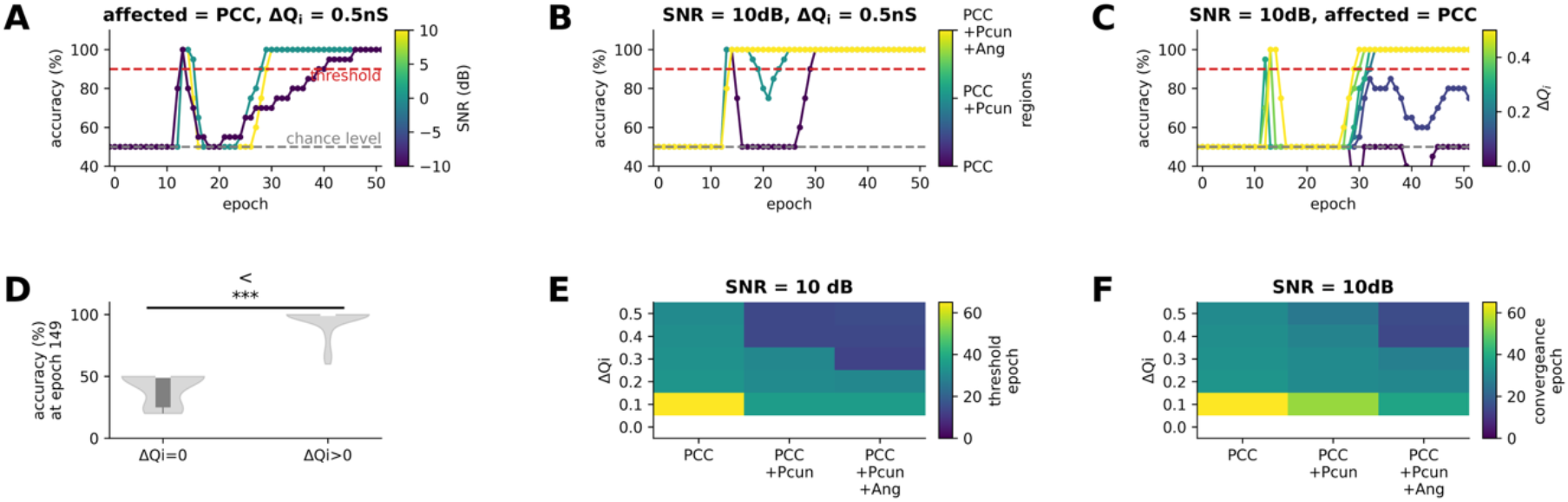
Training and classification performance on whole-brain TVB simulations with a human connectome. (**A-C**) Classification accuracy as a function of training epochs (**A**) for SNRs ranging from −10 dB to 10 dB when only the PCC is affected by E/I imbalance of severity ΔQi = 0.5nS (**B**) for only the PCC, the PCC and PCun, and the PCC, PCun, and AnG affected by E/I imbalance of severity ΔQi = 0.5nS with fixed SNR of −10 dB, and (**C**) for E/I imbalance of severity varying from ΔQi = 0nS to ΔQi = 0.5nS in the PCC with fixed SNR of −10. (**D**) Classification accuracy at epoch 149 for E/I imbalance severity of ΔQi = 0nS and ΔQi > 0nS. (**E-F**) Training epoch during which a 90% accuracy threshold is passed (**E**) and convergence epoch (**F**) as a function of regions affected and E/I imbalance severity for an SNR of 10 dB. *** P < 0.001.

**Figure S3.**
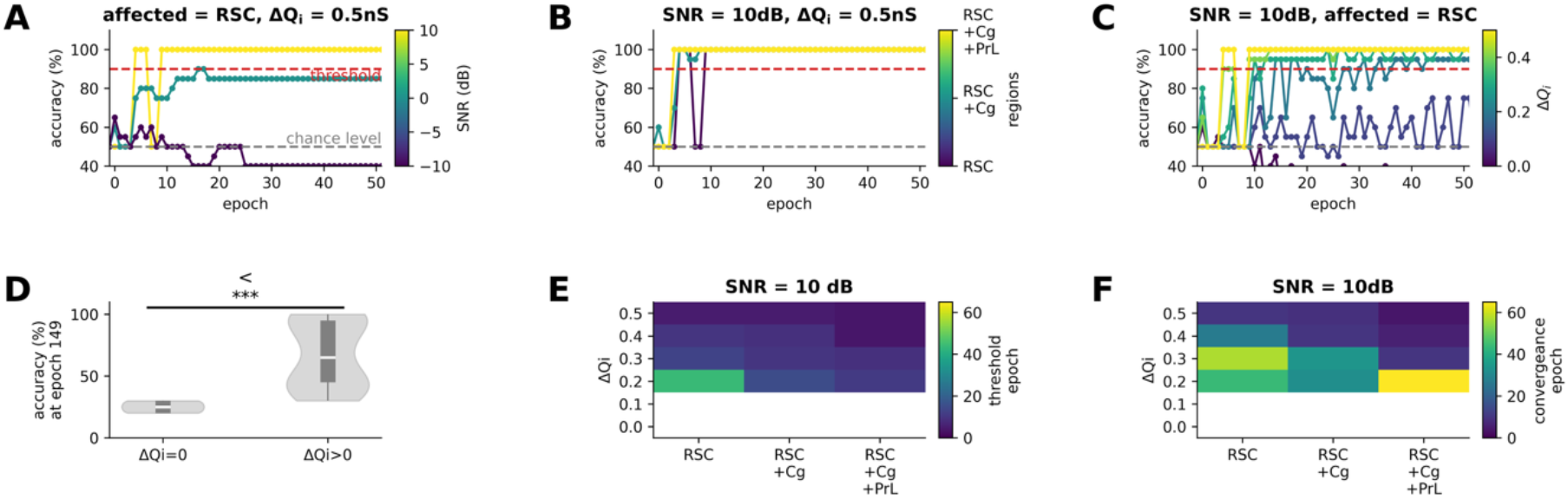
Training and classification performance on whole-brain TVB simulations with a mouse connectome. (**A-C**) Classification accuracy as a function of training epochs (**A**) for SNRs ranging from −10 dB to 10 dB when only the RSC is affected by E/I imbalance of severity ΔQi = 0.5nS (**B**) for only the RSC, the RSC and Cg, and the RSC, Cg, and PrL affected by E/I imbalance of severity ΔQi = 0.5nS with fixed SNR of −10 dB, and (**C**) for E/I imbalance of severity varying from ΔQi = 0nS to ΔQi = 0.5nS in the PCC with fixed SNR of −10. (**D**) Classification accuracy at epoch 149 for E/I imbalance severity of ΔQi = 0nS and ΔQi > 0nS. (**E-F**) Training epoch during which a 90% accuracy threshold is passed (**E**) and convergence epoch (**F**) as a function of regions affected and E/I imbalance severity for an SNR of 10 dB. *** P<0.001.

**Figure S4.**
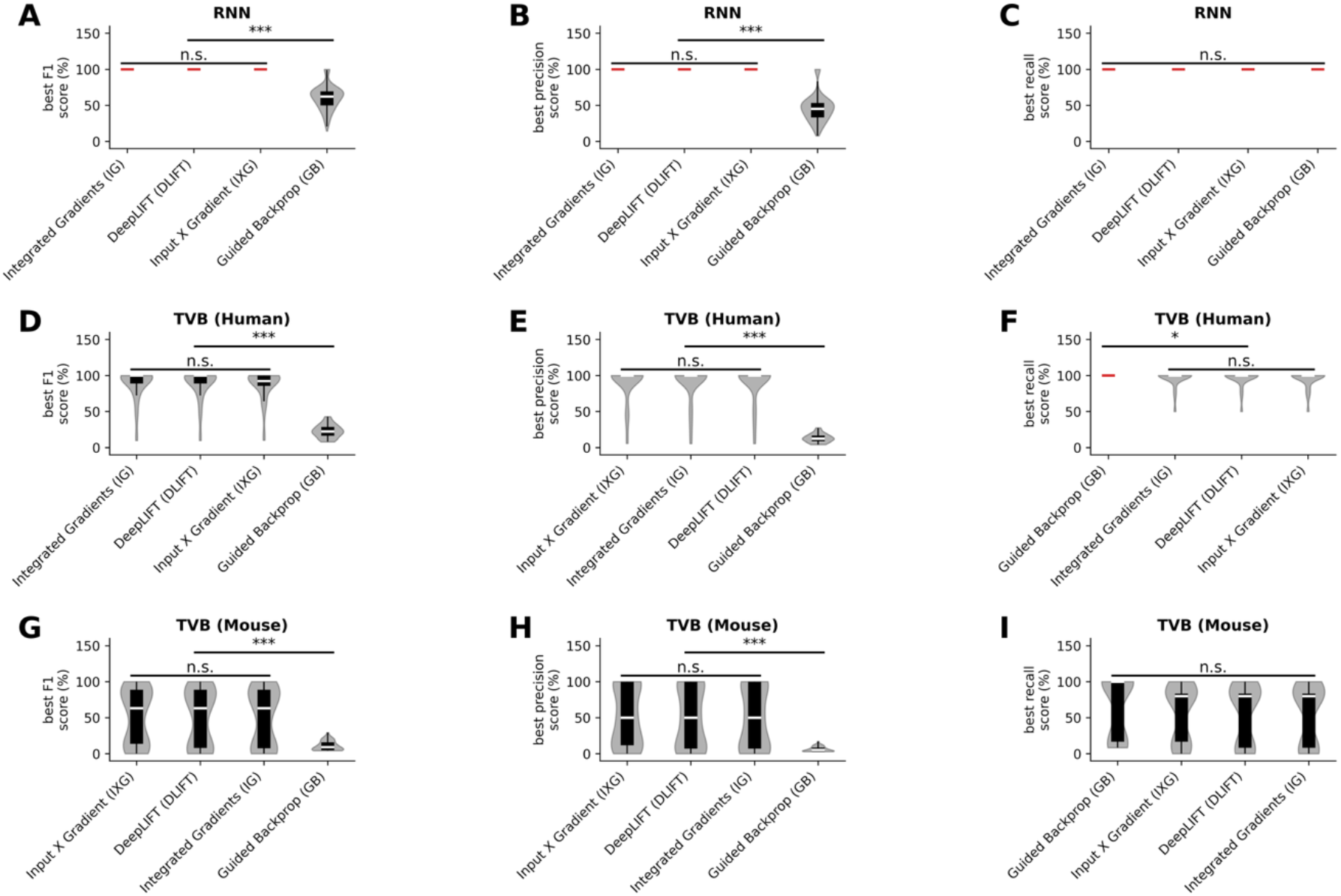
Performance of four different feature attribution methods in detecting affected ROIs. Classifier accuracy quantified by F1score (**A, D, G**), precision (**B, E, H**), and recall (**C, F, I**) in RNN simulations (**A-C**) as well as TVB simulations with human (**D**-**F**) and mouse (**G**-**I**) connectomes across feature attribution methods IXG, DLIFT, IG, and GB. * P < 0.05, *** P < 0.001.

**Figure S5.**
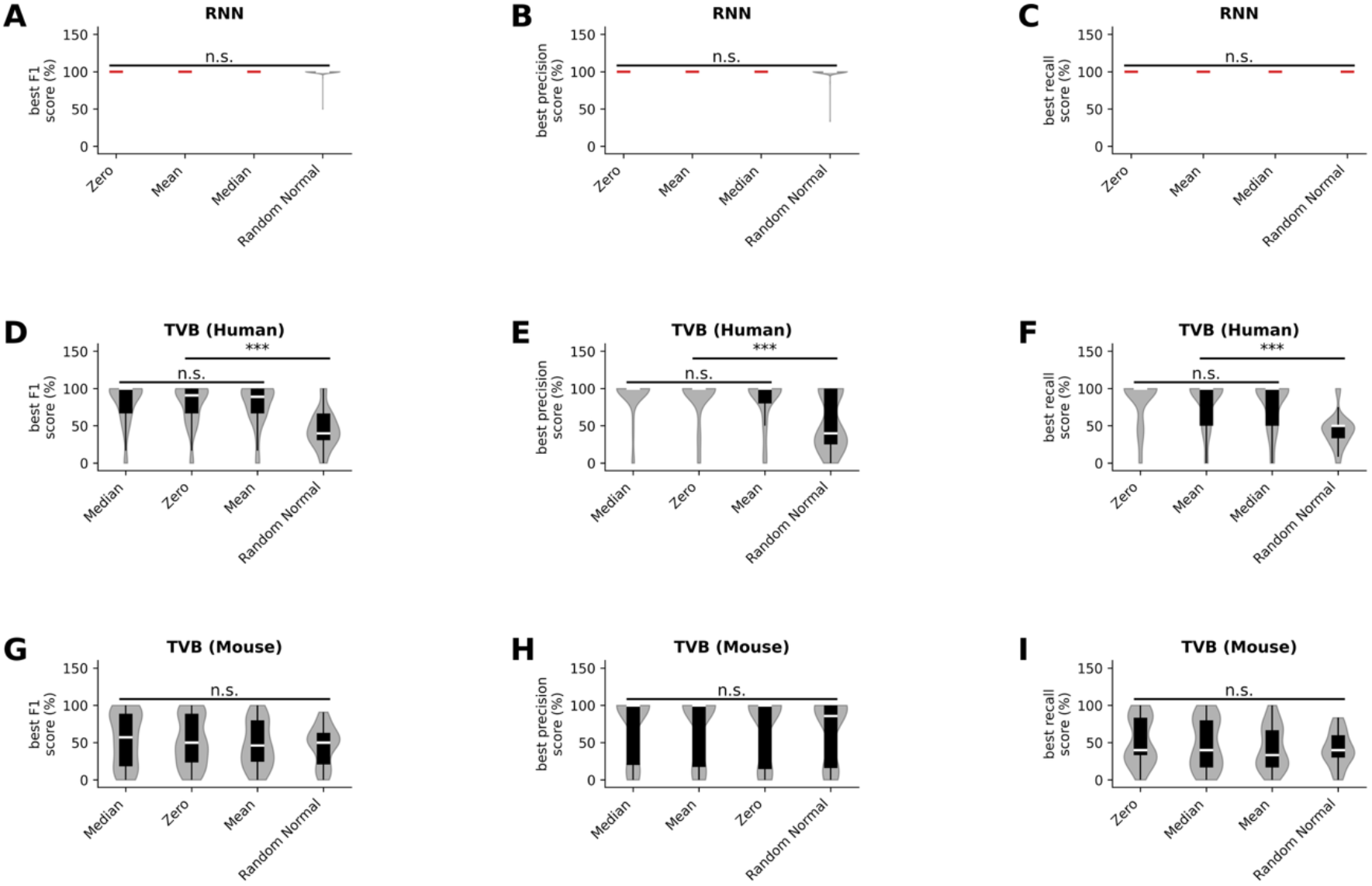
Performance of IG methods with four different baselines in detecting affected ROIs. Classifier accuracy quantified by F1score (**A, D, G**), precision (**B, E, H**), and recall (**C, F, I**) in RNN simulations (**A-C**) as well as TVB simulations with human (**D-F**) and mouse (**G-I**) connectomes across median, mean, zero, and random normal baselines. *** P < 0.001

## Supplementary Tables

**Table S1.**
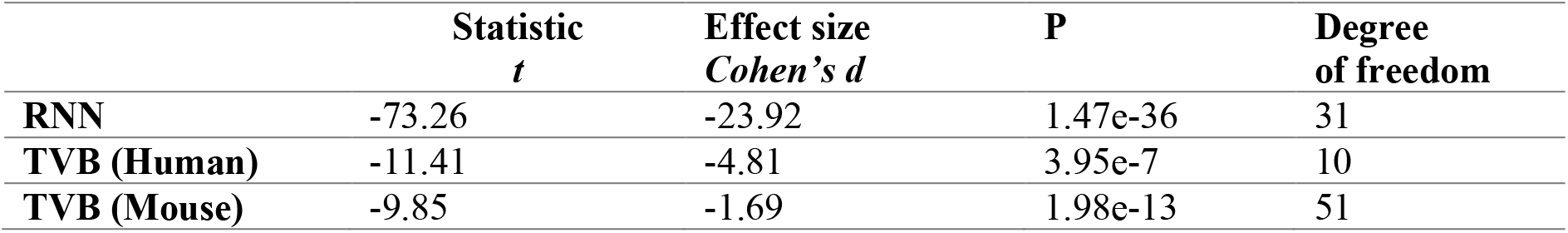
Comparison of models prediction with vs without E/I imbalance. Corresponding to the test performed in the Figure S1D, Figure S2D and Figure S3D.

**Table S2.**
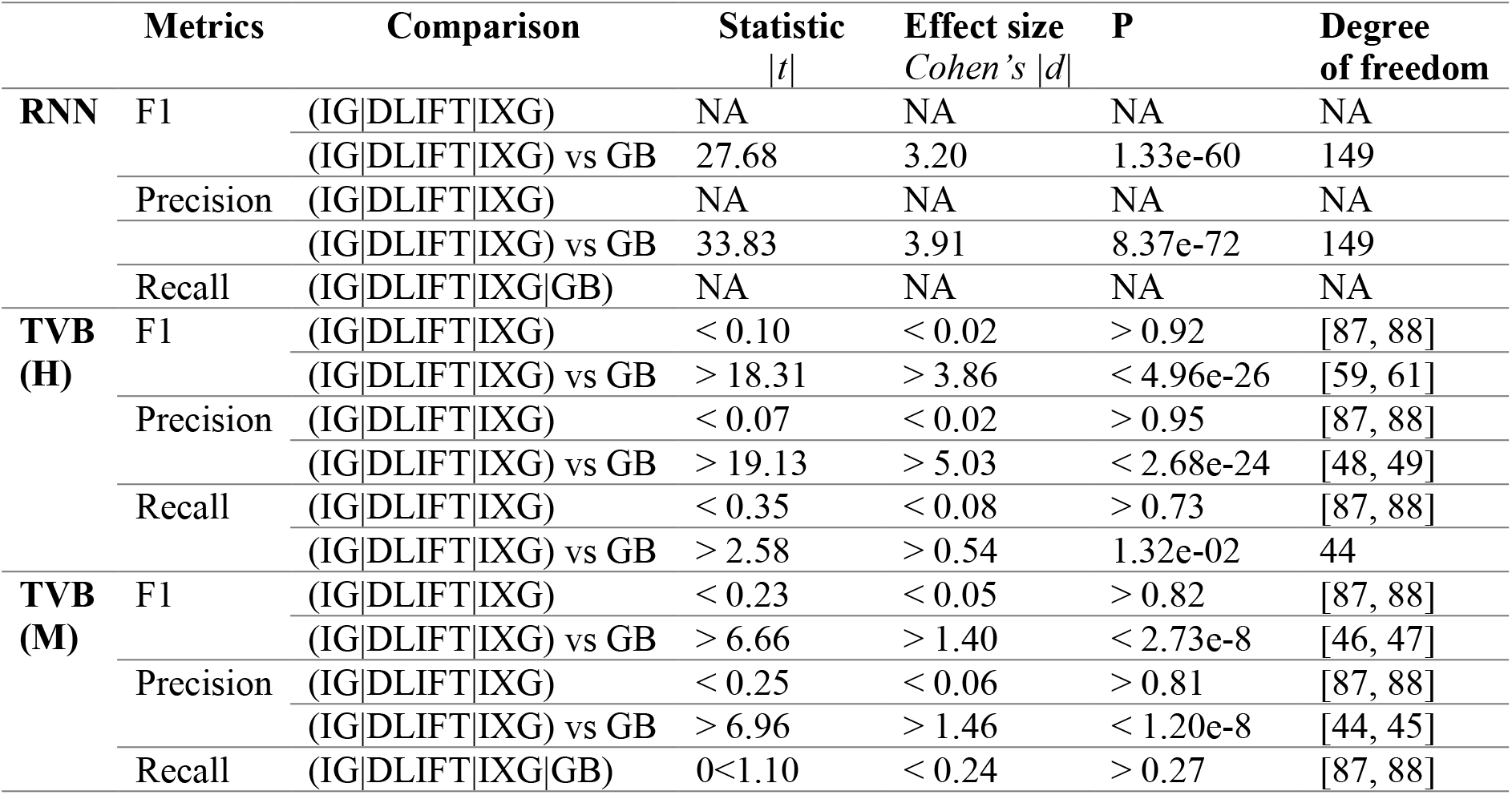
Comparison of attribution methods performance. Corresponding to the test performed in the Figure S4. NA: Not applicable, comparison of constants (i.e. zero variance) that are equal.

**Table S3.** Comparison of IG baseline performance. Corresponding to the test performed in the Figure S5. NA: Not applicable, comparison of constants (i.e. zero variance) that are equal.

|  | Metrics | Comparison | Statistic<br>*t* | Effect size<br>*Cohen's d* | P | Degree<br>of freedom |
| --- | --- | --- | --- | --- | --- | --- |
| **RNN** | F1 | (Z\|MN\|MD) | NA | NA | NA | NA |
|  |  | (Z\|MN\|MD) vs RN | 1.14 | 0.13 | 0.26 | 149 |
|  | Precision | (Z\|MN\|MD) | NA | NA | NA | NA |
|  |  | (Z\|MN\|MD) vs RN | 1.19 | 0.14 | 0.24 | 149 |
|  | Recall | (Z\|MN\|MD\|RN) | NA | NA | NA | NA |
| **TVB**<br>**(H)** | F1 | (Z\|MN\|MD) | < 0.60 | < 0.13 | > 0.55 | [87, 88] |
|  |  | (Z\|MN\|MD) vs RN | > 5.61 | > 1.18 | < 2.29e-7 | [86, 87] |
|  | Precision | (Z\|MN\|MD) | < 0.96 | < 0.03 | > 0.34 | [86, 88] |
|  |  | (Z\|MN\|MD) vs RN | > 4.13 | > 0.87 | < 8.41e-5 | [81, 86] |
|  | Recall | (Z\|MN\|MD) | < 0.87 | < 0.19 | > 0.38 | [87, 88] |
|  |  | (Z\|MN\|MD) vs RN | > 5.78 | > 1.21 | < 1.36e-7 | [80, 82] |
| **TVB**<br>**(M)** | F1 | (Z\|MN\|MD\|RN) | < 1.28 | < 0.27 | > 0.20 | [80, 88] |
|  | Precision | (Z\|MN\|MD\|RN) | < 0.57 | < 0.12 | > 0.57 | [87, 88] |
|  | Recall | (Z\|MN\|MD\|RN) | < 1.94 | < 0.41 | > 0.05 | [77, 88] |

## References

1. Durstewitz D, Koppe G, Meyer-Lindenberg A. Deep neural networks in psychiatry. Mol Psychiatry. 2019;24(11):1583–98.

2. Kriegeskorte N, Golan T. Neural network models and deep learning. Curr Biol. 2019;29(7):R231–R6.

3. Supekar K, de Los Angeles C, Ryali S, Kushan L, Schleifer C, Repetto G, et al. Robust and replicable functional brain signatures of 22q11.2 deletion syndrome and associated psychosis: a deep neural network-based multi-cohort study. Mol Psychiatry. 2024;29(10):2951–66.

4. Supekar K, Ryali S, Yuan R, Kumar D, de Los Angeles C, Menon V. Robust, Generalizable, and Interpretable Artificial Intelligence-Derived Brain Fingerprints of Autism and Social Communication Symptom Severity. Biol Psychiatry. 2022;92(8):643–53.

5. Vieira S, Pinaya WH, Mechelli A. Using deep learning to investigate the neuroimaging correlates of psychiatric and neurological disorders: Methods and applications. Neurosci Biobehav Rev. 2017;74(Pt A):58–75.

6. Breakspear M. Dynamic models of large-scale brain activity. Nat Neurosci. 2017;20(3):340–52.

7. Foss-Feig JH, Adkinson BD, Ji JL, Yang G, Srihari VH, McPartland JC, et al. Searching for Cross-Diagnostic Convergence: Neural Mechanisms Governing Excitation and Inhibition Balance in Schizophrenia and Autism Spectrum Disorders. Biol Psychiatry. 2017;81(10):848–61.

8. Sohal VS, Rubenstein JLR. Excitation-inhibition balance as a framework for investigating mechanisms in neuropsychiatric disorders. Mol Psychiatr. 2019;24(9):1248–57.

9. Gao R, Penzes P. Common mechanisms of excitatory and inhibitory imbalance in schizophrenia and autism spectrum disorders. Current molecular medicine. 2015;15(2):146–67.

10. Tang X, Jaenisch R, Sur M. The role of GABAergic signalling in neurodevelopmental disorders. Nat Rev Neurosci. 2021;22(5):290–307.

11. Lányi O, Koleszár B, Schulze Wenning A, Balogh D, Engh MA, Horváth AA, et al. Excitation/inhibition imbalance in schizophrenia: a meta-analysis of inhibitory and excitatory TMS-EMG paradigms. Schizophrenia. 2024;10(1):56.

12. Nelson SB, Valakh V. Excitatory/Inhibitory Balance and Circuit Homeostasis in Autism Spectrum Disorders. Neuron. 2015;87(4):684–98.

13. Rubenstein JL, Merzenich MM. Model of autism: increased ratio of excitation/inhibition in key neural systems. Genes Brain Behav. 2003;2(5):255–67.

14. Hollestein V, Poelmans G, Forde NJ, Beckmann CF, Ecker C, Mann C, et al. Excitatory/inhibitory imbalance in autism: the role of glutamate and GABA gene-sets in symptoms and cortical brain structure. Transl Psychiatry. 2023;13(1):18.

15. Horder J, Petrinovic MM, Mendez MA, Bruns A, Takumi T, Spooren W, et al. Glutamate and GABA in autism spectrum disorder-a translational magnetic resonance spectroscopy study in man and rodent models. Transl Psychiatry. 2018;8(1):106.

16. Padmanabhan A, Lynch CJ, Schaer M, Menon V. The Default Mode Network in Autism. Biol Psychiatry Cogn Neurosci Neuroimaging. 2017;2(6):476–86.

17. Hu ML, Zong XF, Mann JJ, Zheng JJ, Liao YH, Li ZC, et al. A Review of the Functional and Anatomical Default Mode Network in Schizophrenia. Neurosci Bull. 2017;33(1):73–84.

18. Menon V. 20 years of the default mode network: A review and synthesis. Neuron. 2023;111(16):2469–87.

19. Gadgil S, Zhao Q, Pfefferbaum A, Sullivan EV, Adeli E, Pohl KM. Spatio-Temporal Graph Convolution for Resting-State fMRI Analysis. Med Image Comput Comput Assist Interv. 2020;12267:528–38.

20. Ngo GH, Khosla M, Jamison K, Kuceyeski A, Sabuncu MR. Predicting individual task contrasts from resting-state functional connectivity using a surface-based convolutional network. Neuroimage. 2022;248:118849.

21. Zhang Y, Farrugia N, Bellec P. Deep learning models of cognitive processes constrained by human brain connectomes. Med Image Anal. 2022;80:102507.

22. Lei D, Qin K, Pinaya WHL, Young J, Van Amelsvoort T, Marcelis M, et al. Graph Convolutional Networks Reveal Network-Level Functional Dysconnectivity in Schizophrenia. Schizophr Bull. 2022;48(4):881–92.

23. Wang L, Li K, Hu XP. Graph convolutional network for fMRI analysis based on connectivity neighborhood. Netw Neurosci. 2021;5(1):83–95.

24. Ryali S, Supekar K, Chen T, Kochalka J, Cai W, Nicholas J, et al. Temporal Dynamics and Developmental Maturation of Salience, Default and Central-Executive Network Interactions Revealed by Variational Bayes Hidden Markov Modeling. PLoS Comput Biol. 2016;12(12):e1005138.

25. Lurie DJ, Kessler D, Bassett DS, Betzel RF, Breakspear M, Kheilholz S, et al. Questions and controversies in the study of time-varying functional connectivity in resting fMRI. Network Neuroscience. 2020;4(1):30–69.

26. Calhoun VD, Miller R, Pearlson G, Adali T. The Chronnectome: Time-Varying Connectivity Networks as the Next Frontier in fMRI Data Discovery. Neuron. 2014;84(2):262–74.

27. Ryali S, Zhang Y, de Los Angeles C, Supekar K, Menon V. Deep learning models reveal replicable, generalizable, and behaviorally relevant sex differences in human functional brain organization. Proc Natl Acad Sci U S A. 2024;121(9):e2310012121.

28. Eitel F, Schulz MA, Seiler M, Walter H, Ritter K. Promises and pitfalls of deep neural networks in neuroimaging-based psychiatric research. Exp Neurol. 2021;339.

29. Ghassemi M, Oakden-Rayner L, Beam AL. The false hope of current approaches to explainable artificial intelligence in health care. Lancet Digit Health. 2021;3(11):e745–e50.

30. Sanz-Leon P, Knock SA, Woodman MM, Domide L, Mersmann J, McIntosh AR, et al. The Virtual Brain: a simulator of primate brain network dynamics. Front Neuroinform. 2013;7:10.

31. Glasser MF, Smith SM, Marcus DS, Andersson JLR, Auerbach EJ, Behrens TEJ, et al. The Human Connectome Project’s neuroimaging approach. Nature Neuroscience. 2016;19(9):1175–87.

32. Oh SW, Harris JA, Ng L, Winslow B, Cain N, Mihalas S, et al. A mesoscale connectome of the mouse brain. Nature. 2014;508(7495):207-+.

33. Uhlhaas PJ, Singer W. Neuronal dynamics and neuropsychiatric disorders: toward a translational paradigm for dysfunctional large-scale networks. Neuron. 2012;75(6):963–80.

34. Sundararajan M, Taly A, Yan QQ. Axiomatic Attribution for Deep Networks. Pr Mach Learn Res. 2017;70.

35. Shrikumar A, Greenside P, Kundaje A. Learning Important Features Through Propagating Activation Differences. Pr Mach Learn Res. 2017;70.

36. Kindermans P-J, Hooker S, Adebayo J, Alber M, Schütt KT, Dähne S, et al. The (un) reliability of saliency methods. Explainable AI: Interpreting, explaining and visualizing deep learning. 2019:267–80.

37. Springenberg J, Dosovitskiy A, Brox T, Riedmiller M, editors. Striving for Simplicity: The All Convolutional Net. ICLR (workshop track); 2015.

38. Gevaert A, Rousseau AJ, Becker T, Valkenborg D, De Bie T, Saeys Y. Evaluating feature attribution methods in the image domain. Mach Learn. 2024;113(9):6019–64.

39. Desikan RS, Ségonne F, Fischl B, Quinn BT, Dickerson BC, Blacker D, et al. An automated labeling system for subdividing the human cerebral cortex on MRI scans into gyral based regions of interest. Neuroimage. 2006;31(3):968–80.

40. Supekar K, Uddin LQ, Khouzam A, Phillips J, Gaillard WD, Kenworthy LE, et al. Brain Hyperconnectivity in Children with Autism and its Links to Social Deficits. Cell Rep. 2013;5(3):738–47.

41. Watts DJ, Strogatz SH. Collective dynamics of ‘small-world’networks. Nature. 1998;393(6684):440–2.

42. Ryali S, Supekar K, Chen T, Menon V. Multivariate dynamical systems models for estimating causal interactions in fMRI. Neuroimage. 2011;54(2):807–23.

43. Sanz-Leon P, Knock SA, Spiegler A, Jirsa VK. Mathematical framework for large-scale brain network modeling in The Virtual Brain. Neuroimage. 2015;111:385–430.

44. Goldman JS, Kusch L, Aquilue D, Yalcinkaya BH, Depannemaecker D, Ancourt K, et al. A comprehensive neural simulation of slow-wave sleep and highly responsive wakefulness dynamics. Front Comput Neurosc. 2023;16.

45. Volo MD, Romagnoni A, Capone C, Destexhe A. Biologically Realistic Mean-Field Models of Conductance-Based Networks of Spiking Neurons with Adaptation. Neural Comput. 2019;31(4):653–80.

46. Klein A, Tourville J. 101 labeled brain images and a consistent human cortical labeling protocol. Front Neurosci-Switz. 2012;6.

